# Affinity-matured CD72-targeting Nanobody CAR T-cells Enhance Elimination of Antigen-Low B-cell Malignancies

**DOI:** 10.1101/2025.05.09.653155

**Authors:** Adila Izgutdina, Tasfia Rashid, William C. Temple, Bonell Patiño-Escobar, Sujata Walunj, Huimin Geng, Hiroyuki Takamatsu, Daniel Gil-Alós, Amrik S. Kang, Emilio Ramos, Szu-Ying Chen, Haley Johnson, Matthew A. Nix, Akul Naik, Constance M. Yuan, Hao-Wei Wang, Sarah Aminov, Srabani Sahu, Rebecca C. Larson, Christopher Carpenter, Fernando Salangsang, Paul Phojanakong, Juan Antonio Camara Serrano, Isa Tariq, Ons Zakraoui, Veronica Steri, Antonio Valeri, Joaquin Martinez-Lopez, Marcela V. Maus, Samir Parekh, Amit Verma, Nirali N. Shah, Arun P. Wiita

## Abstract

**Background:** Chimeric antigen receptor (CAR) T-cell therapies are highly efficacious for several different hematologic cancers. However, for most CAR T targets it is observed that low surface antigen density on tumors can significantly reduce therapeutic efficacy. Here, we explore this dynamic in the context of CD72, a surface antigen we recently found as a promising target for refractory B-cell cancers, but for which CD72 low antigen density can lead to therapeutic resistance in preclinical models.

**Methods:** Primary samples were accessed via institutional review board-approved protocols. Affinity-matured and humanized nanobody clones were previously described in Temple et al.^1^ CAR T-cells were generated via lentiviral transduction. *In vitro* cytotoxicity assays were performed using luciferase-labeled cell lines. *In vivo* studies were performed using cell line- or patient-derived xenografts implanted in NOD *scid* gamma (NSG) mice.

**Results:** We first confirmed ubiquitous CD72 expression across a range of primary B-cell non-Hodgkin lymphomas. We further found that after resistance to CD19-directed therapies, across both B-cell acute lymphoblastic leukemia (B-ALL) models and primary tumor samples, surface CD72 expression was largely preserved while CD22 expression was significantly diminished. Affinity maturation of a nanobody targeting CD72, when incorporated into chimeric antigen receptor (CAR) T-cells, led to more effective elimination *in vitro* of isogenic models of CD72 low-expressing tumors. These results suggested that nanobody-based CAR T-cells (nanoCARs) may exhibit a similar relationship between binder affinity, antigen expression, and efficacy as previously demonstrated only for scFv-based CAR T-cells. Surprisingly, however, this significantly improved *in vitro* efficacy only translated to modest *in vivo* survival benefit. As a parallel strategy to enhance CAR T function, we found that the small molecule bryostatin could also significantly increase CD72 surface antigen density on B-cell malignancy models. Structural modeling and biochemical analysis identified critical residues improving CD72 antigen recognition of our lead affinity-matured nanobody.

**Conclusions:** Together, these findings support affinity-matured CD72 nanoCARs as a potential immunotherapy product for CD19-refractory B-cell cancers. Our results also suggest that for B-ALL in particular, CD72 may be a preferable second-line immunotherapy target over CD22.

**What is already known on this topic:** Previous work using single chain variable fragment (scFv) based CAR Ts has suggested that improving affinity for target antigen could potentially help mitigate tumor resistance mediated by antigen downregulation, or baseline low antigen density. However, it is unknown whether this same dynamic holds for CAR T-cells that utilize different antigen recognition elements, such as nanobodies.

**What this study adds:** Here we show that affinity maturation of nanobody-based CAR T-cells (nanoCARs) targeting CD72 can improve their *in vitro* efficacy versus CD72-low tumors; however, *in vivo* efficacy differences are more modest. Furthermore, we show that for refractory B-cell malignancies, surface CD72 appears preserved after CD19 resistance even in situations where CD22 is strongly downregulated.

**How this study might affect research, practice or policy:** CD72 warrants further investigation as a preferred immunotherapy target in the context of CD19-refractory B-cell cancers, though nanobody affinity maturation is not a universal solution to the challenge of low tumor surface antigen density.

## INTRODUCTION

Chimeric antigen receptor (CAR) T-cells are a powerful therapeutic strategy for patients with refractory B-cell cancers. However, while there are four different CD19-targeting CAR T-cells currently approved by the U.S. Food and Drug Administration, many patients unfortunately still relapse and need new options. Of many potential mechanisms of resistance to CAR T therapy^2^, one that has been recognized in diffuse large B-cell lymphoma (DLBCL) is decreased density of CD19 antigen at the tumor cell surface^3^. In B-ALL, loss of the CAR-targeting epitope on CD19 (ref.^4^) has also been observed to cause resistance. Similar dynamics of clinical relapse due to decreased antigen density have been defined for other CAR T targets such as CD22 in B-cell acute lymphoblastic leukemia (B-ALL)^5^ and B-cell maturation antigen (BCMA) in multiple myeloma^6^. Other targets have shown similar post-treatment downregulation or diminished efficacy in tumors with baseline low antigen density^7^. Therefore, an important design challenge in CAR T-cells is how to more effectively capture and eliminate tumors with low surface density of the target of interest.

One potential strategy to address low antigen density tumors is to target a different antigen altogether. In the case of B-cell cancers, several alternative CAR T targets have been proposed, such as CD20, CD22, CD79a/b, CD37, CD70, and others^8^ for B-cell non-Hodgkin lymphomas (B-NHL), or CD22 for B-cell acute lymphoblastic leukemia (B-ALL)^5,9^. In the latter case, however, despite CD22 CAR Ts leading to impressive clinical responses in refractory B-ALL patients, remissions have not been durable^5,10^. New options are thus needed for refractory B-ALL patients in particular.

To address this challenge, we have recently identified CD72 as a promising antigen that appears widely expressed in both B-ALL and B-NHL^1,11^. In normal tissues CD72 is largely specific for B-cells with minimal expression on other cell types^11^. In prior work, we previously developed proof-of-principle CAR T-cells against CD72 using a synthetically designed nanobody display library^11^, then further optimized these “nanoCARs” for clinical translation using a framework humanization strategy^1^. While this approach has shown encouraging results, we have also noted in preclinical models that tumors with lower antigen density may evade elimination by our previously described anti-CD72 nanoCARs^1,11^.

The most common antigen recognition element used in CAR T-cells are antibody fragments called single chain variable fragments, or scFv’s. Prior work has shown that engineering scFv’s to achieve higher affinity for the target antigen can eliminate tumor cells with lower antigen density^12–15^, though others have found that increased affinity may not always be advantageous in the context of low antigen density^16^. Nanobodies, derived from heavy chain-only antibodies, are half the size of scFv’s and can engage different binding modes with target antigens^17^. Despite the rising excitement around nanobody-based CAR T-cells, including one commercially-approved product^18^ and numerous others in (pre)clinical development^19,20,21^, it remains unknown if increasing nanobody affinity can improve CAR T-cell efficacy versus antigen-low tumors.

Here, we test this hypothesis, finding that affinity maturation of nanobody-based CAR T-cells can indeed lead to enhanced elimination of CD72-low tumor cells *in vitro*, albeit with more modest impacts *in vivo*. Analysis of patient B-ALL tumors also reveal preserved CD72 expression after CD19 CAR T relapse, even when CD22 surface expression is diminished. We further use computational and biophysical approaches to identify key residues responsible for this improved nanoCAR activity. Finally, we demonstrate a pharmacological approach to increase CD72 antigen density, which could be considered as a combination strategy with CD72 nanoCARs. Taken together, our results show that affinity maturation is a potential strategy to improve nanobody-based CAR T-cell efficacy, while also supporting CD72-targeting nanoCARs as a promising therapeutic option for patients with refractory B-cell malignancies. For B-ALL in particular, our results also suggest that CD72 may be a preferable second-line CAR T target after CD19.

## MATERIALS and METHODS

### CAR T binders and vector constructs

Nanobody affinity maturation was described in Temple et al.^1^ Humanization of framework region of anti-CD72 nanobody clones NbD4.7 and NbD4.13 was performed by substituting additional framework mutations present in the “H24” nanobody (H24 sequence described in Temple et al.^1^). All anti-CD72 nanobody sequences were cloned into the same CAR backbone that included a mutated IgG4 hinge domain^22^, CD28 transmembrane and costimulatory domains, and CD3ζ signaling domain, which we previously found to optimize efficacy of an anti-CD72 nanoCAR design^1^; transduction efficiency was monitored via a fused Green Fluorescent Protein (GFP) marker. As a clinically relevant comparator, CD19 binder (FMC63 scFv) and Empty CAR (CAR backbone only, no binder) were cloned into the tisagenlecleucel backbone, with CD8 hinge and transmembrane domain, 4-1BB costimulatory domain, and CD3ζ signaling domain.

### CAR T-cell production

Primary human T-cells were isolated from anonymous healthy donor leukapheresis product (StemCell Technologies). Cells were enriched for CD4+ and CD8+ populations, using enrichment kits EasySep Human CD4 T Cell Iso Kit (StemCell Technologies, 17952) and EasySep Human CD8 T Cell Iso Kit (StemCell Technologies, 17953) and then viably frozen. CD4+ and CD8+ T-cells were thawed and cultured in OpTimizer medium with CTS supplement (Thermo Scientific, A1048501), with 5% human AB serum (Valley Medical, HP1022), and penicillin/streptomycin. The next day, T-cells were stimulated with 20 μL of CD3/CD28 Dynabeads (Thermo Fisher Scientific, 11131-D) per 1e6 T-cells for 4-5 days. T-cells were cultured in the presence of 10 ng/mL of recombinant interleukin-7 (PeproTech, 200-07) and interleukin-15 (PeproTech, 200-15). T-cells were transduced with CAR lentivirus 1 day post bead stimulation. Transduction efficiency was assessed 3-4 days after transduction. As CAR constructs all include a fused GFP tag, the percentage of GFP positive cells by flow cytometry determined the percentage of CAR positive cells.

### Murine studies

Murine studies were conducted at the UCSF Helen Diller Family Comprehensive Cancer Center Preclinical Therapeutics Core in accordance with UCSF Institutional Animal Care and Use Committee guidelines (Approval Number: AN194778-01). 6-9 week old NSG mice of mixed male and female sex (NOD.Cg-*Prkdcscid Il2rgtm1Wjl*/SzJ, Jackson Laboratories) were used for all experiments.

### Statistical analysis

Statistical analysis of data was performed in GraphPad Prism Version 10.1.1 (270). Unpaired student *t*-test was performed to compare two groups and one way ANOVA with Tukey’s correction was used to compare one independent variable between more than two groups. Log-rank Mantel-Cox test was done to evaluate survival in mouse studies. Data points are presented as mean +/- standard deviation, unless stated otherwise. *p*-values <0.05 were considered significant (“ns” p>0.05, *p<0.05, **p<0.01, ***p ≤ 0.001, ****p ≤ 0.0001).

Additional Methods are included in the Supplementary Materials file attached to this manuscript.

## RESULTS

### CD72 is widely expressed on B-NHL patient tumors and cell line tumor models

In prior work, we confirmed ubiquitous surface expression of CD72 in a cohort of pediatric B-ALL samples via flow cytometry, as well as showed via immunohistochemistry that 96% of evaluated adult diffuse large B-cell lymphomas (DLBCL) had positive CD72 staining^11^. While these prior results were encouraging for the promise of CD72 as a cellular therapy target for a wide catchment of B-cell cancers, here we sought to extend these findings.

Initial analysis of microarray transcriptome data from Brune et al.^23^ (downloaded from Gene Expression Omnibus accession GSE12453) confirmed *CD72* mRNA expression across normal B-cells and several B-NHL subtypes including DLBCL, follicular lymphoma, and Burkitt lymphoma (**Fig. 1A**). In addition, in DLBCL and chronic lymphocytic leukemia (CLL) cohorts (GSE10846, GSE23967, GSE22762), high expression of *CD72* was associated with significantly reduced overall survival (**Fig. 1B-D**). These transcriptome-level analyses suggest a potential prognostic value of *CD72* expression in B-NHL patient populations.

**Figure 1.**
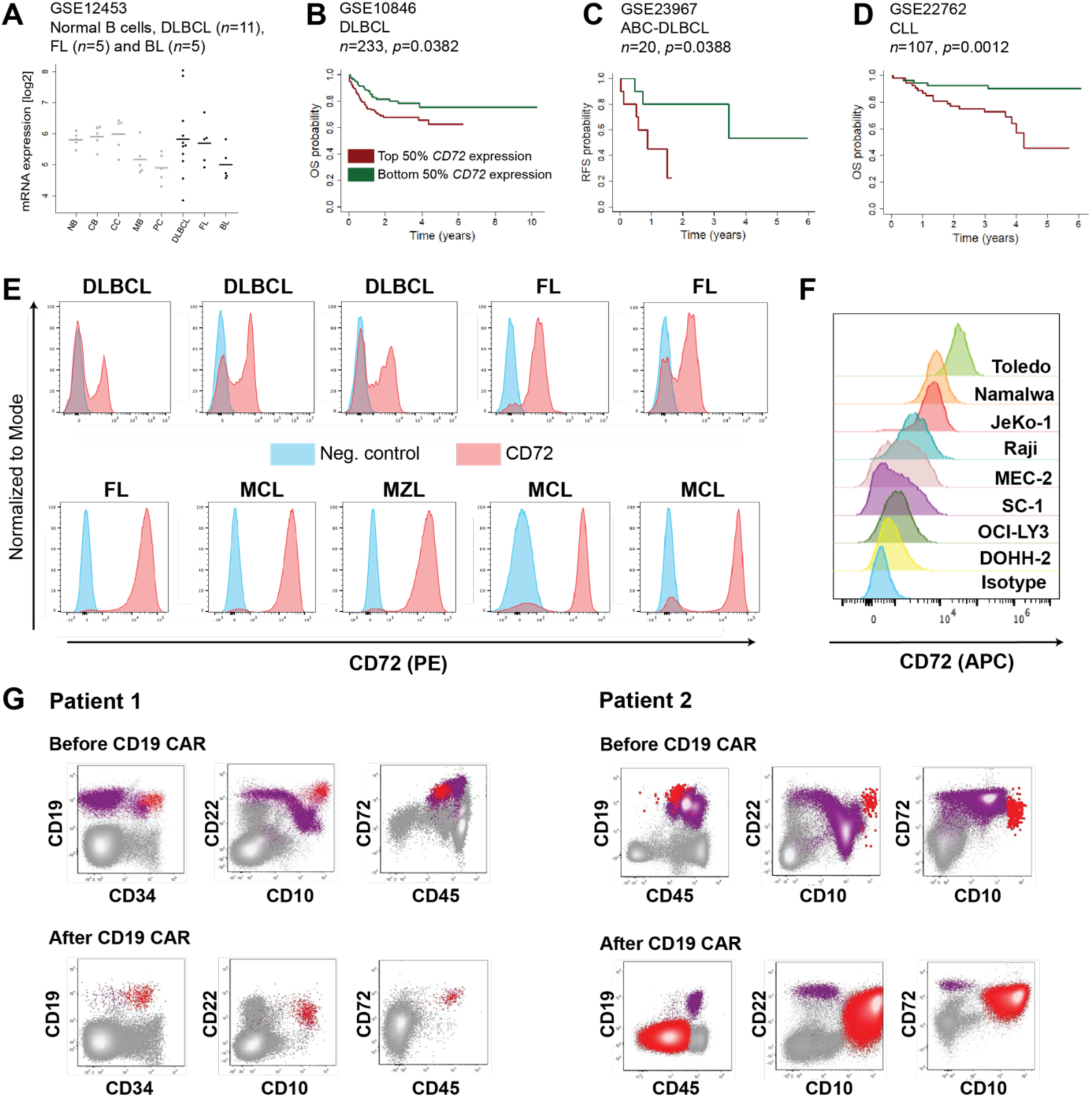
Transcriptome analysis and flow cytometry confirm CD72 is a B-cell specific target ubiquitously present in B-cell malignancies and retained after CD19 CAR T relapse. A) *CD72* transcript expression from microarray profiling in healthy B-cells and B-NHL lymphoma cells. GSE12453: DLBCL (*n*=11), FL (*n*=5) and BL (*n*=5). **B-D)** Kaplan–Meier plots of overall survival (OS) and relapse-free survival (RFS) in B-cell lymphoma patient cohorts selected by *CD72* expression: top 50% expression in red and bottom 50% expression in green. **B)** GSE10846, DLBCL, treated with R-CHOP, *n*=233, *p*=0.0382 by Log-Rank test. **C)** GSE23967, ABC-DLBCL, treated with R-CHOP, *n*=20, *p*=0.0388 by Log-Rank test. **D)** GSE22762, CLL, *n*=107, *p*=0.00124 by Log-Rank test. **E)** Flow cytometry evaluation of CD72 expression in primary B-NHL lymphoma patient tumor samples from Kanazawa University, Japan (top panel, isotype control was used as negative control) and Hospital Universitario 12 de Octubre, Madrid, Spain (bottom panel, fluorescence minus one was used as negative control). **F)** Flow cytometry of B-NHL lymphoma tumor cell lines with a broad range of CD72 expression. **G)** Flow cytometry of two pediatric B-ALL patients profiled for expression of CD19, CD22 and CD72 before and after CD19 CAR T therapy relapse. B-ALL blasts CD10+/CD45 low+ (Patient 1) and CD10+/CD45 low+/- (Patient 2).

To corroborate microarray expression data, we analyzed CD72 surface expression by flow cytometry in B-NHL primary tumor samples from two independent institutions, as well as commonly used B-NHL cell line models. We profiled nine primary lymphoma samples from Kanazawa University, Japan, including DLBCL (*n* = 4), follicular lymphoma (FL) (*n* = 2), mantle cell lymphoma (MCL) (*n* = 2) and Burkitt lymphoma (BL) (*n* = 1). CD72 was uniformly expressed on tumor cells, with mean fluorescence intensity (MFI) fold-change compared to isotype control ranging from 27-fold to 149-fold, similar to that observed for CD19 (**Fig. 1E, Supp. Fig. 1A-B**). In addition, we profiled eleven tumors from Hospital Universitario 12 de Octubre, Madrid, Spain, which included FL (*n* = 1), MCL (*n* = 5), marginal zone lymphoma (*n* = 4), and lymphoplasmacytic lymphoma (*n* =1). CD72 was again present on all tumors at high levels, with MFI increase from 78 to 1118-fold difference compared to fluorescence minus one negative control (**Fig. 1E, Supp. Fig. 1A**). Complementary to the primary samples, analysis of 8 commonly used B-NHL cell lines showed a range of CD72 expression; on these cell lines, using the same antibody clone (3F3) but different fluorophore conjugate (APC, vs. PE in primary sample staining above), we found CD72 typically showed lower MFI than CD19 (**Fig. 1F**, **Supp. Fig. 1C**).

We further investigated CD72 expression in the context of resistance to the first-line immunotherapy target CD19. We recently showed in two *in vitro* models of CD19 immunotherapy resistance that surface CD19 was downregulated, mimicking the dynamic seen in some patients^24^. Surprisingly, in these same models CD22 was simultaneously downregulated despite not being under any direct therapeutic pressure^24^. Here, we also evaluated CD72 expression in these same models. Intriguingly, in the REH cell line model CD72 showed less downregulation than CD22, both by flow cytometry for surface protein (<1-fold decrease for CD72 vs. >8-fold reduction for CD22) as well as at the transcript level by re-analysis of our previously published single cell RNA-seq data (**Supp. Fig. 1D-E**). To extend these findings from model systems, we analyzed CD72 surface expression on two pediatric B-ALL patient tumor samples that showed CD19 antigen loss post-CD19 CAR T-relapse. In these samples, CD22 surface expression was strongly downregulated, whereas CD72 surface expression was preserved (**Fig. 1G**). Taken together, these findings suggest that CD72 may be a more suitable second line target than CD22 after CD19-based therapy relapse, and further suggest that CD72 is a targetable antigen across a range of adult B-cell lymphomas.

### Affinity matured CD72 CAR demonstrates largely equivalent efficacy to prior humanized H24 CAR in CD72 high-expressing models

We next set out to further optimize our CD72 nanoCARs, with the goal of improving efficacy versus B-NHL across a range of antigen densities as well as to avoid antigen-low relapse in B-ALL or B-NHL. For other targets, one strategy proposed to achieve these goals is to increase the affinity of the antigen binding domain included in the CAR^12^. However, while this approach has proven successful for some scFv-based CAR Ts, it is unknown whether it could also be successful for nanobody-based CAR Ts.

To investigate this possibility, we took advantage of a series of affinity matured anti-CD72 nanobody variants that we have previously described^1^ (**Fig. 2A**). Briefly, we previously compared the *in vitro* anti-tumor efficacy of several of these affinity-matured clones (“NbD4.1”, “NbD4.3”, “NbD4.7”, “NbD4.13”; designed using overlapping degenerate PCR from the parental “NbD4” clone), versus the humanized “H24” nanobody clone, with each incorporated into a 2^nd^ generation CAR backbone with a CD28-based costimulatory domain^1^. In that work we also confirmed superiority of the specific H24 (non-affinity-matured) sequence to several other humanized variants as well as the parental (fully llama backbone) “NbD4” clone^1^. Biolayer interferometry confirmed that recombinant Fc domain fusion of either the NbD4.7 and NbD4.13 affinity-matured nanobody showed a K_D_ of ∼1.8 and ∼0.8 nM, respectively, compared to 33 nM for H24^1^. However, in that prior work, when we tested the resulting CAR Ts in a 24 hour co-culture assay with high CD72-expressing cell lines SEM (B-ALL model) or JeKo-1 (MCL model), we found that affinity maturation did not improve *in vitro* cytotoxicity compared to H24-based CAR Ts^1^.

**Figure 2.**
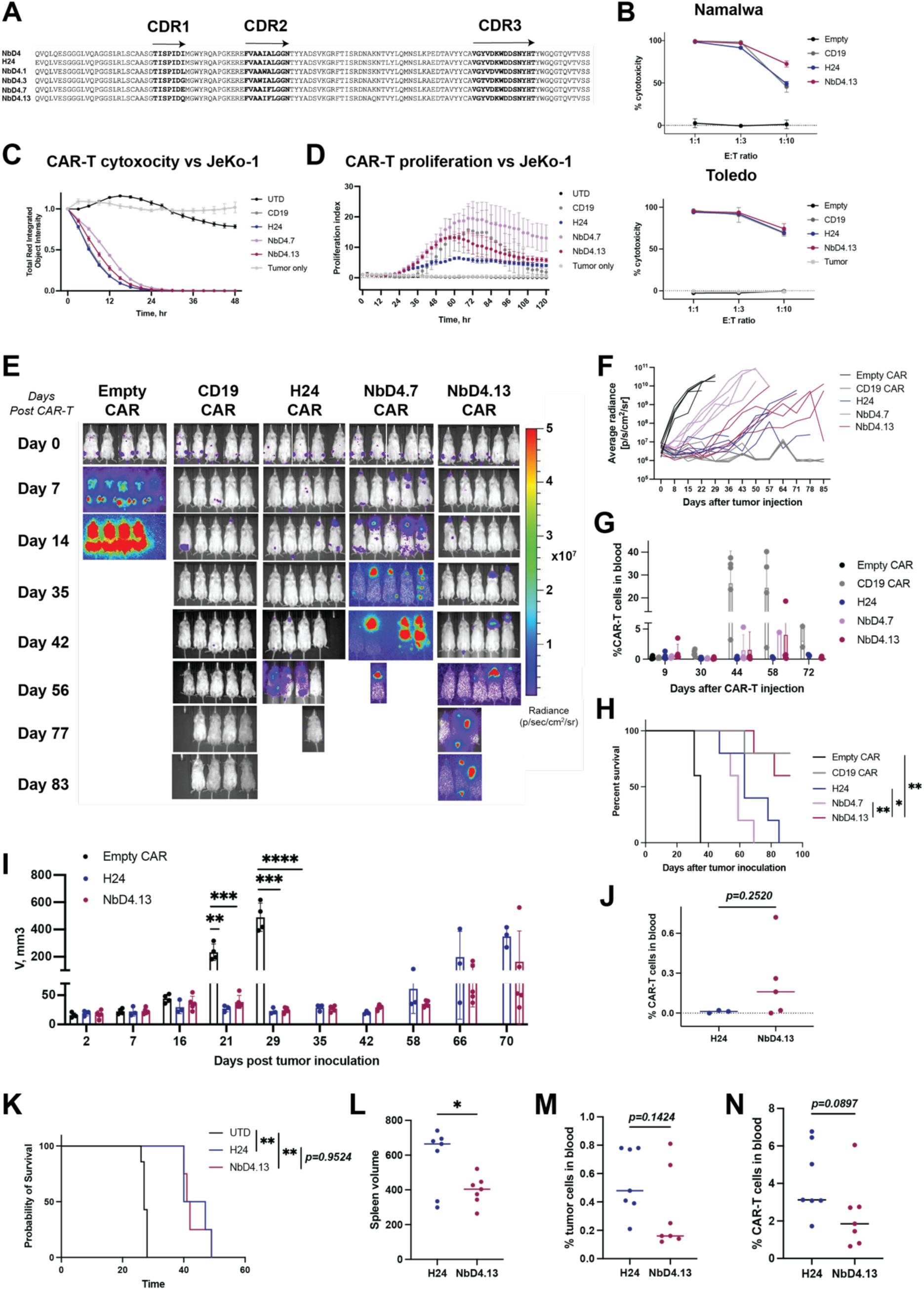
Affinity matured CD72 CAR-T-cells show increased tumor control *in vitro* and *in vivo* against B-NHL lymphoma models compared to H24 CAR. A) Amino acid sequences of anti-CD72 nanobody used in CAR T designs, including parental NbD4 (originally described in ref.^11^), humanized H24, affinity matured NbD4.1, NbD4.3, NbD4.7, and NbD4.13 (originally described in ref.^1^). Complementarity determining regions (CDRs) highlighted. B) *In vitro* 48-hour luciferase-based cytotoxicity assay of CD72 CAR Ts versus lymphoma cell lines Namalwa and Toledo. Data normalized to untransduced T-cells. *n*=3 technical replicates, performed at noted E:T ratios. **C-D)** Kinetics of CD72 CAR-T cytotoxicity and proliferation versus mCherry-labeled JeKo-1 MCL cell line by Incucyte live cell imaging. Data normalized to time zero of each well. *n*=3 technical replicates. Performed at 1:3 Effector:Tumor (E:T) ratio. **E-F)** Luciferase-labeled JeKo-1 (1e6 tumor cells) implanted intravenously in NSG mice (*n*=5 per group) and treated with 3.5e6 CAR T-cells 7 days after. Bioluminescence images (**E**) and quantified average radiance (**F**) plot of JeKo-1 tumor in mice treated with Empty CAR (negative control), CD19 CAR, H24, NbD4.7 or NbD4.13 CAR. **G)** Quantification of blood CAR T expansion in murine peripheral blood over time. CAR Ts quantified based on human CD3+/GFP+ cells. Statistical analysis was performed by two-way ANOVA with Tukey’s multiple comparisons test. **H)** Kaplan–Meier curve of overall survival in JeKo-1 study. Statistical analysis was performed by log-rank (Mantel-Cox) test. (\**p*<0.05, \*\**p*<0.01). **I)** Murine study with 1e6 mantle cell lymphoma PDX DFBL44685 cells injected into mice intravenously and subsequently intravenously implanted with 5e6 CAR Ts 10 days after. Murine tumor burden assessed by spleen ultrasound at 7 day intervals. **J)** CAR T expansion in murine blood on day 51 after PDX tumor injection. **K)** Survival analysis of high tumor burden stress test with 2e6 PDX cells implanted intravenously and treated with CAR-Ts 14 days after. Kaplan–Meier curve of overall survival. Statistical analysis was performed by log-rank (Mantel-Cox) test. **L)** Spleen volume, **M)** tumor burden in blood and **N)** CAR T percentage in blood at 6 weeks after tumor injection. Statistical analysis in I, J, L-N) was performed by Student’s *t*-test. (ns *p*>0.05, \**p*<0.05, \*\**p*<0.01, \*\*\**p*≤0.001, ****p≤0.0001). Independent T-cell donors were used for studies in (**B**), (**C-D**), (**F-H**), and (**I-N**).

In the study here, consistent with our prior published results, we also found largely equivalent *in vitro* cytotoxicity of affinity-matured CARs and H24-based CAR against two additional CD72 high-expressing B-NHL models, Toledo (DLBCL model) and Namalwa (BL model) (**Fig. 2B**). We further evaluated the kinetics of anti-tumor cytotoxicity of affinity matured CAR variants, H24, and a CD19-targeting positive control CAR (tisagenlecleucel mimic) versus the JeKo-1 cell line using an Incucyte^®^ Live-Cell imaging platform. Consistent with the single time-point assays above, the Incucyte assay demonstrated equivalent cytotoxicity of all CAR designs at 24 hours of exposure (**Fig. 2C**). However, we found superior expansion of affinity matured CAR Ts compared to H24 (**Fig. 2D**).

We next investigated *in vivo* performance of these various CD72 nanoCAR designs in a model of JeKo-1 cells intravenously implanted in NSG mice. Surprisingly, the NbD4.7 design showed very poor tumor control, with all mice rapidly relapsing (**Fig. 2E**), despite having very similar affinity for CD72 compared to NbD4.13. Intriguingly, by bioluminescence imaging we found that despite some NbD4.13-treated mice showing evidence of earlier tumor outgrowth than H24-treated mice, ultimately, NbD4.13-treated mice showed significantly improved survival compared to H24 (*p* = 0.035 by log-rank test) (**Fig. 2F-G**). There was no appreciable difference in peripheral blood CAR T expansion kinetics for any anti-CD72 CAR design, though the positive control anti-CD19 CAR did show increased expansion and improved tumor control (**Fig. 2H, Supp. Fig. 2A**). Taken together, these findings indicate that increasing nanobody affinity for CD72 can lead to widely varying impacts on tumor control, either greatly diminishing anti-tumor efficacy (NbD4.7) or potentially improving it (NbD4.13). Notably, in this case, these dramatically different *in vivo* outcomes are mediated by a change of just 2 amino acids between these two nanobody clones (**Fig. 2A**).

### NbD4.13 leads to modestly improved tumor control over H24 in a PDX MCL model

Given the above findings, for the remainder of our work we focused on the NbD4.13 clone as the best-performing affinity-matured anti-CD72 nanobody variant. To further investigate *in vivo* performance of NbD4.13, we established a patient-derived xenograft (PDX) model of mantle cell lymphoma (MCL) (obtained from ProXe Biobank^25^) that showed strong CD72 expression (**Supp. Fig. 2B)**. In an initial study, we injected 1e6 tumor cells and administered 5e6 CAR T-cells (NbD4.13, H24, or negative control) at Day 10. In NbD4.13-treated vs. H24-treated mice we observed modestly reduced tumor burden, as measured by spleen ultrasound, as well as increased peripheral blood CAR T-cells, as measured by flow cytometry (**Fig. 2I-J, Supp. Fig. 2C**). Hypothesizing that impacts of our affinity matured nanoCARs may be more prominent with greater initial tumor burden, we performed a subsequent study using a higher dose of tumor cells (2e6). In this study, at 6 weeks we did observe significantly decreased tumor burden in NbD4.13-treated mice vs. H24, although overall survival was similar (**Fig. 2K-M**). Taken together, these findings in a CD72 high-expressing PDX model also suggest modestly improved *in vivo* performance of NbD4.13 vs. H24-based CAR Ts.

### Affinity matured NbD4.13 CD72 CAR is superior to humanized H24 CAR versus an isogenic “antigen low” model *in vitro*

We next sought to test our initial hypothesis, that an affinity-matured CAR T would show the greatest advantage in anti-tumor efficacy in the context of low CD72 expression. In our prior work^1^ we found that H24-treated JeKo-1 cells relapsed *in vivo* with ∼3.3-fold lower CD72 antigen density than untreated JeKo-1 cells. We isolated tumor cells from relapsed murine spleens and expanded these cells *ex vivo* to create a “CD72lo^r^” model (r = relapsed). In an Incucyte live cell-imaging assay, we observed significantly faster JeKo-1 CD72lo^r^ tumor elimination for affinity-matured CAR variants compared to H24 (NbD4.13 vs H24 *p*=0.039; NbD4.7 vs H24 *p*=0.067 by *t*-test) (**Fig. 3A**).

**Figure 3.**
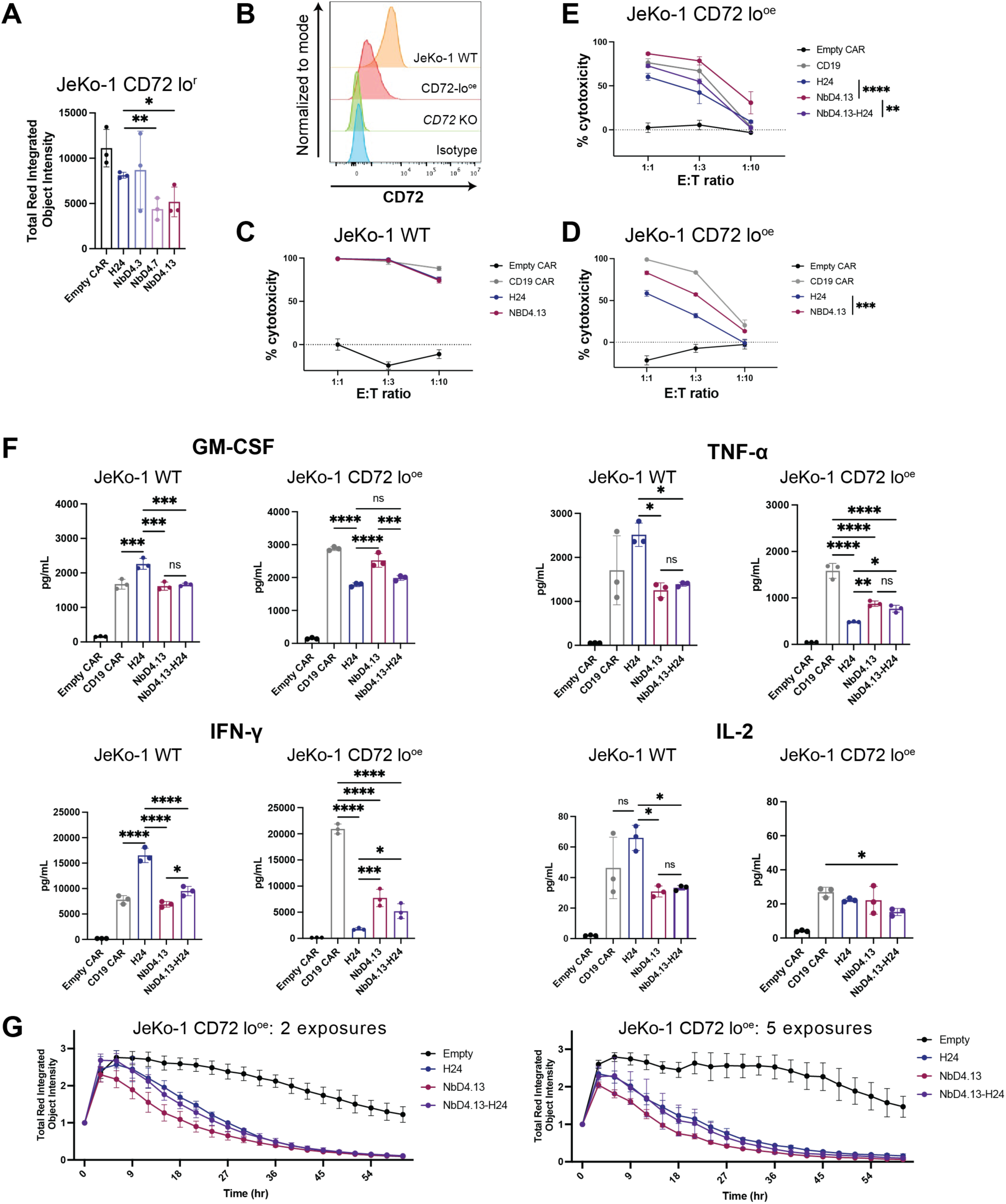
Affinity matured CAR-Ts have superior effector function to H24 *in vitro* versus Jeko1-CD72 low model. **A)** JeKo-1 tumor cells were isolated from mice implanted with WT tumor and relapsed after H24 CAR therapy (from our prior study^1^; previously shown to have decreased CD72 expression at relapse). These mCherry-labeled JeKo-1 CD72lo^r^ cells were co-cultured with CAR-Ts at 1:3 E:T in Incucyte live cell analyzer. The bar graph represents cytotoxicity at 36 hours of co-culture. Data analyzed by unpaired *t*-test. **B)** Flow cytometry plots showing CD72 expression in JeKo-1 WT, JeKo-1 *CD72* KO and engineered JeKo-1 CD72-lo^oe^ model. **C-D)** *In vitro* 24-hour luciferase-based cytotoxicity assay of Empty, CD19, H24, NbD4.13 CAR Ts versus JeKo-1 WT (**C**) and JeKo-1 CD72 lo^oe^ (**D**) models. Normalized to tumor cells, *n*=3 technical replicates. **E)** *In vitro* 24-hour luciferase-based cytotoxicity assay of Empty, CD19, H24, NbD4.13 and NbD4.13-H24 CAR-Ts versus JeKo-1 CD72lo^oe^ model. Normalized to untransduced T-cells, *n*=3 technical replicates. Statistical analysis in D) and E) was performed by two-way ANOVA with Tukey’s multiple comparisons test (\*\**p*<0.01, \*\*\**p*≤0.001, \*\*\*\**p*≤0.0001). Performed with CAR T-cells generated from an independent donor from (**C-D**). **F)** Cytokine profiling of CD72 CAR T supernatant by Luminex assay (see Methods) after 24hr exposure to JeKo-1 WT and JeKo1-lo^oe^ tumor models at 1:1 E:T. Statistical analysis of data is performed by ordinary one-way ANOVA (ns *p*>0.05, \**p*<0.05, \*\**p*<0.01, \*\*\**p* ≤ 0.001, \*\*\*\**p* ≤ 0.0001). **G)** Cytotoxicity kinetics of affinity matured (NbD4.13), affinity matured+humanized (NbD4.13-H24), and humanized (H24) CD72 CAR-Ts with repetitive stimulation with JeKo-1 CD72 lo^oe^ tumor (mCherry-labeled), 2 and 5 exposures to tumor. Data obtained from Incucyte live cell analyzer. Data normalized to time zero of each well. *n*=3 technical replicates. Performed with CAR T-cells generated from an independent donor from (**C-F**).

We also developed a second isogenic model of low CD72 expression. We used CRISPR/Cas9 to knock out *CD72* in JeKo-1 cells and then exogenously re-expressed the CD72 open reading frame via stable lentiviral transduction under a SFFV promoter. This “CD72lo^oe^” (oe = overexpression) model resulted in surface CD72 expression ∼4.7x lower than the parental JeKo-1, based on MFI by flow cytometry (**Fig. 3B**). In this model we also observed significantly improved anti-tumor efficacy of the NbD4.13 affinity-matured variant compared to H24 (**Fig. 3C-E**).

We first characterized additional CAR T phenotypes when cultured with JeKo-1 WT cells. We found that the NbD4.13 variant showed increased central memory and decreased effector phenotype compared to H24 variant, based on CD62L/CD45RA staining (**Supp. Fig. 3A**). We also found significantly decreased LAG-3 staining for NbD4.13 compared to H24, though expression of other exhaustion markers (PD-1, TIM-3) was similar (**Supp. Fig. 3B**). In the same assay, NbD4.13 also showed increased T-cell activation (CD69) and degranulation (CD107a) compared to H24 (**Supp. Fig. 3C**).

We then compared findings between these CAR Ts cultured with either JeKo-1 WT or CD72lo^oe^ models. Against the WT JeKo-1 cell line, H24 nanoCARs showed increased secretion of cytokines associated with CAR activity including tumor necrosis factor α (TNF-α), granulocyte-monocyte colony stimulating factor (GM-CSF), interleukin-2 (IL-2) and interferon ψ (IFN-ψ), compared to NbD4.13 (**Fig. 3F**). However, versus CD72lo^oe^ model, both NbD4.13 and a hybrid humanized + affinity matured variant (“NbD4.13-H24”, combining point mutations found in H24 or NbD4.13 clones (see sequence in **Supp. Fig. 2D**)) showed increased or equivalent secretion of the above cytokines compared to H24 (**Fig. 3F**). Several other cytokines profiled did not show significant differences (**Supp. Fig. 3D-E**). Given recent evidence that CAR T-cell avidity for target cells may be an important determinant of anti-tumor efficacy^26^, we also performed acoustic force spectroscopy using the Lumicks platform. We found evidence of significantly increased avidity against both JeKo-1 WT and JeKo-1 CD72lo^oe^ cells for NbD14.13 compared to H24 (**Supp. Fig. 3F**).

A common *in vitro* approach used to simulate *in vivo* CAR T dynamics of multiple target cell encounter is the “repetitive stimulation” assay^27^. In this assay, we cocultured the same CAR Ts with fresh CD72lo^oe^ cells every 3 days for up to 5 co-cultures. In an Incucyte assay, NbD4.13- based CARs consistently showed more rapid tumor elimination, after both 2 and 5 repetitive stimulations, compared to H24 (**Fig. 3G**). Taken together, these findings demonstrate that affinity maturation of the anti-CD72 nanobody in our CAR construct can drive improved anti-tumor efficacy *in vitro* versus CD72-low expressing tumor models.

### Affinity matured CD72 CAR only marginally improves on H24 *in vivo* against antigen low JeKo-1 model

Given our *in vitro* findings above versus the CD72lo^oe^ JeKo-1 model, we further sought to evaluate whether this same improved anti-tumor efficacy of affinity-matured CARs would be observed *in vivo*. We intravenously implanted CD72lo^oe^ JeKo-1 cells in NSG mice and treated 1 week later with CAR Ts (**Fig. 4A**). Indeed, in line with our initial hypothesis, we observed improved initial tumor control for the affinity-matured NbD4.13 and hybrid NbD4.13-H24 CARs compared to H24 up to Day 28 of the study (**Fig. 4B**), though CAR T expansion was not different (**Fig. 4C**). Surprisingly, however, the CD72lo^oe^ JeKo-1 model exhibited a more aggressive phenotype *in vivo* when compared to WT JeKo-1, leading to rapid relapse in all treated mice. In this context, we were only able to resolve a modest improvement in overall survival for NbD4.13-H24 CARs vs. H24 (*p* = 0.044 by log-rank test) and a trend toward improved survival for NbD4.13 CARs vs. H24 (*p* = 0.164) (**Fig. 4D**). Single cell RNA-seq analysis of spleens from mice post-relapse indicated no detectable CAR T-cells and minimal differences between H24, NbD4.13, and Empty (negative control) treated tumor (**Supp. Fig. 4**).

**Figure 4.**
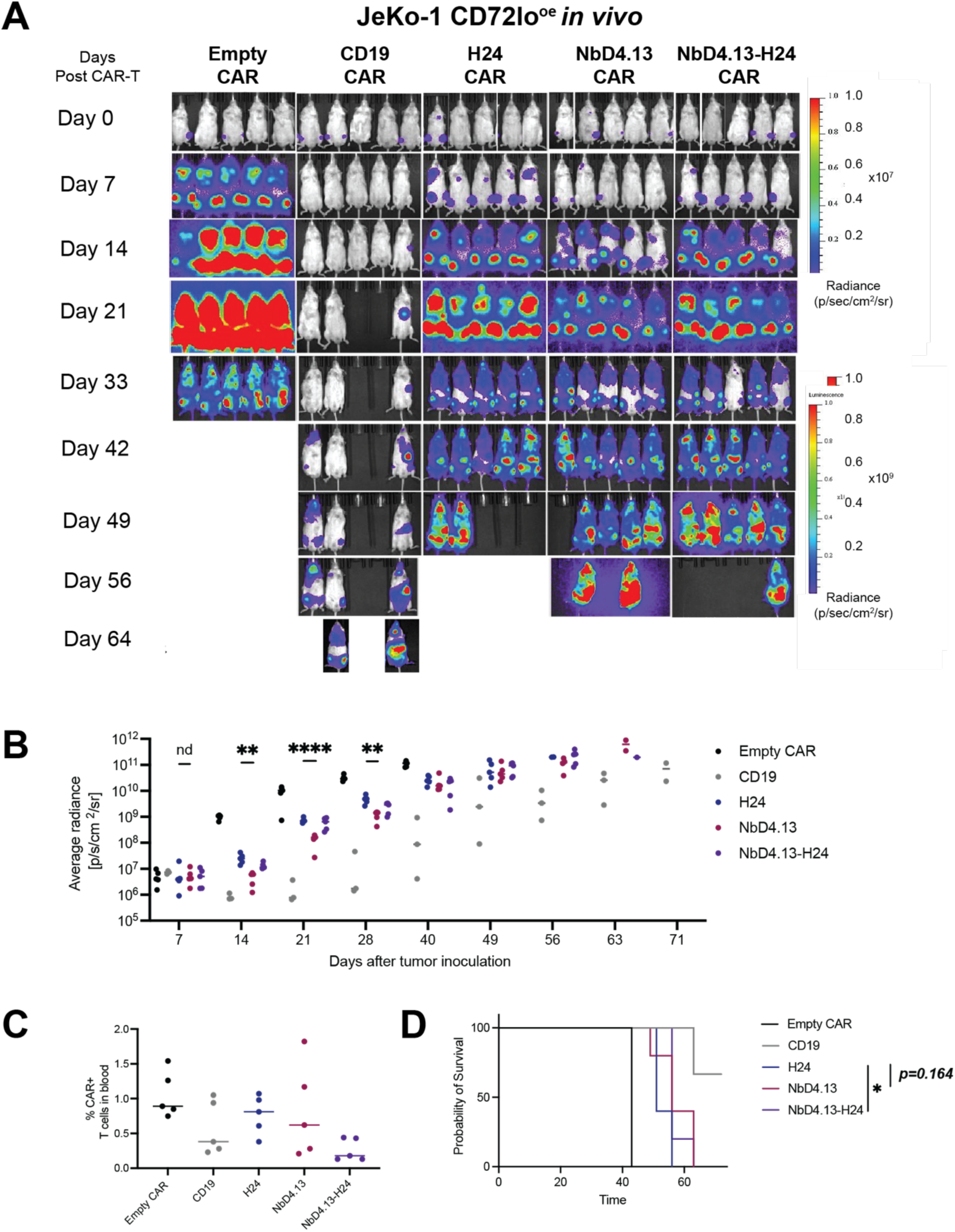
Affinity matured and humanized CAR-Ts have a modest survival benefit versus JeKo-1 low model. NSG mice were implanted with 0.5e6 JeKo-1 CD72 lo^oe^ tumor cells through tail vein injection and treated with 5e6 CAR-T-cells 7 days after (*n*=5 per arm). **A)** Tumor burden assessed by weekly bioluminescence imaging. **B)** Quantified BLI data over time. Data analyzed by multiple unpaired *t*-test. (ns *p*>0.05, \*\**p*<0.01, \*\*\*\**p*≤0.0001). **C)** Peripheral CAR-T expansion on day 9 after CAR-T injection. Statistical analysis by ordinary one-way ANOVA. **D)** Kaplan–Meier curve of overall survival. Statistical analysis was performed by log-rank (Mantel-Cox) test (ns *p*>0.05, \**p*<0.05).

Taken together, these findings indicate that while affinity-matured nanoCARs led to clearly improved *in vitro* anti-tumor activity versus CD72lo tumors, markedly enhanced *in vivo* efficacy is not achieved as readily.

### Bryostatin pharmacologically upregulates surface CD72 and enhances CD72 nanoCAR anti-tumor efficacy

In parallel with the affinity maturation studies above, we investigated whether pharmacologically increasing tumor surface CD72 antigen levels could be a complementary strategy to enhance function of our nanoCARs. To this end, we noted that the Protein Kinase C inhibitor bryostatin was previously reported to increase surface CD22 expression in B-cell cancer models and drive improved anti-tumor activity of anti-CD22 CAR T-cells^16^. We thus hypothesized that similar downstream signaling driven by bryostatin may also increase surface CD72. Using two B-NHL models that natively express low CD72 (SC-1 and DOHH-2 (**Fig. 1F**)) as well as JeKo-1 WT and SEM cells (**Supp. Fig. 5A**), we found that treatment with 1 nM bryostatin over 72 hr led to significant surface upregulation of not only CD22, consistent with the prior study^16^, but also significant upregulation of CD72 and CD19 (**Fig. 5A** and **Supp. Fig. 5A**). When we cocultured different CD72 nanoCARs with SC-1 cells after treating with bryostatin (1 nM, 72 hr), we measured a higher degree of tumor elimination at lower effector to target cell ratios (**Fig. 5B**). These findings were consistent with the increase in tumor surface CD72 enhancing CAR T efficacy.

**Figure 5.**
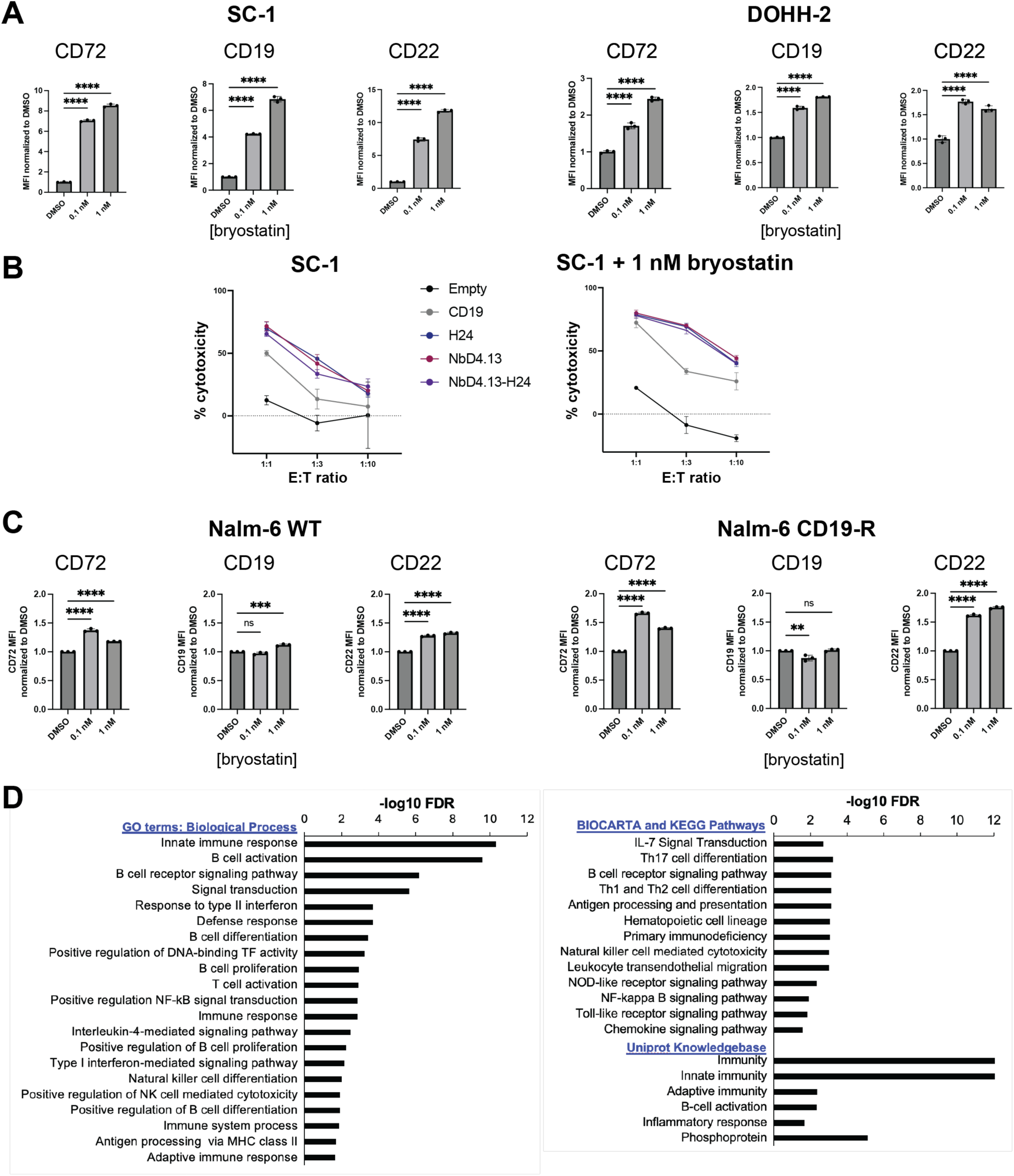
Bryostatin is an antigen modulator of B-cell associated targets. **A)** SC-1 or DOHH-2 cells treated with 0.1% DMSO or increasing concentrations of bryostatin. Surface antigen density of CD72, CD19, CD22 profiled by flow cytometry and normalized to DMSO treated cells. **B)** SC-1 untreated cells and cells pre-treated with 1 nM of bryostain for 72 hr were then co-cultured with CAR Ts in a 24 hr luciferase-based cytotoxicity assay at 1:1, 1:3 and 1:10 E:T ratios. **C)** Nalm-6 WT and CD19 therapy relapsed-models (Nalm-6 CD19-R) were treated with 0.1% DMSO or increasing concentrations of bryostatin. Surface antigen density of CD72, CD19, and CD22 profiled by flow cytometry and normalized to DMSO treated cells. Statistical analysis in A) and C) was performed by ordinary one-way ANOVA with Dunnett’s multiple comparisons test (ns *p*>0.05, \**p*<0.05, \*\**p*<0.01, \*\*\**p* ≤ 0.001, \*\*\*\**p* ≤ 0.0001). **D)** Pathway analysis of bulk RNA-seq of bryostatin treated SC-1 cells demonstrates 591 upregulated genes and 204 downregulated genes (FDR<0.05). B-cell activation and signaling are among the top pathways represented among upregulated genes.

As an alternate pharmacologic strategy, we also investigated the DNA methyltransferase inhibitor azacytidine, which we and others have found can lead to surface upregulation of other cancer immunotherapy targets^28,29^. Azacytidine could also modestly increase CD72 antigen density (**Supp. Fig. 5B**). We further investigated whether bryostatin can also increase surface CD72 in models of relapse after CD19-directed immunotherapy. Indeed, using the NALM-6 model shown in **Supp. Fig. 1E**, bryostatin treatment (0.1 or 1 nM, 72 hr) could increase surface expression of CD72. Interestingly, bryostatin also partially restored expression of CD22 in this model, though with less impact on CD19 (**Fig. 5C**).

Given the ability of bryostatin to simultaneously increase antigen density of multiple B-cell-specific surface markers, we sought to further investigate this small molecule’s mechanistic impact on B-ALL tumor cells. We thus treated SC-1 cells for 72 hr with 1 nM bryostatin and performed bulk RNA-seq (**Supp. Fig. 5C-E**). Pathway analysis of upregulated transcripts in bryostatin-treated cells compared to vehicle-treated cells showed highly significant upregulation of B-cell specific signaling mechanisms including B-cell differentiation and activation, B-cell receptor signaling, positive regulation of NF-κB signal transduction, chemokine signaling and inflammatory response pathways (**Fig. 5D**).

Taken together, these findings indicate that bryostatin co-treatment could be a promising future strategy to combine with affinity matured CD72-targeting nanoCARs to enhance anti-tumor efficacy on CD72-low tumors, and this small molecule acts by strongly activating B-cell-specific signaling pathways.

### Structural modeling and investigation of the CD72-nanobody interface

Our results above suggest that a small number of mutations in our nanobody constructs can strongly modulate their affinity for target antigens. We further sought to mechanistically identify the most critical residues mediating the interaction between CD72 and either the originally-described NbD4 clone^11^, the subsequently humanized H24 clone^1^, or the affinity matured NbD4.13 clone. We first used AlphaFold^30^ and HADDOCK^31^ to predict the structure of each of nanobody in complex with the human CD72 extracellular domain (ECD). This structural modeling suggested that, as expected given their common origin from the NbD4 clone, these nanobodies share a similar epitope on the CD72 ECD. We predicted this epitope to be present at the junction of the coiled coil and lectin-like globular domain of CD72, between amino acids 210 and 250 (**Fig. 6A**). We specifically examined the NbD4.13 interface with CD72, which suggested that the single mutation in CDR2, Ala52Phe (A52F), may have the most critical role in improving affinity compared to the NbD4 parental clone. This alteration appears to create a favorable interaction between an aromatic ring of the nanobody Phe52 with the proline ring Pro222 and adjacent phenylalanine Phe223 on CD72.

**Figure 6.**
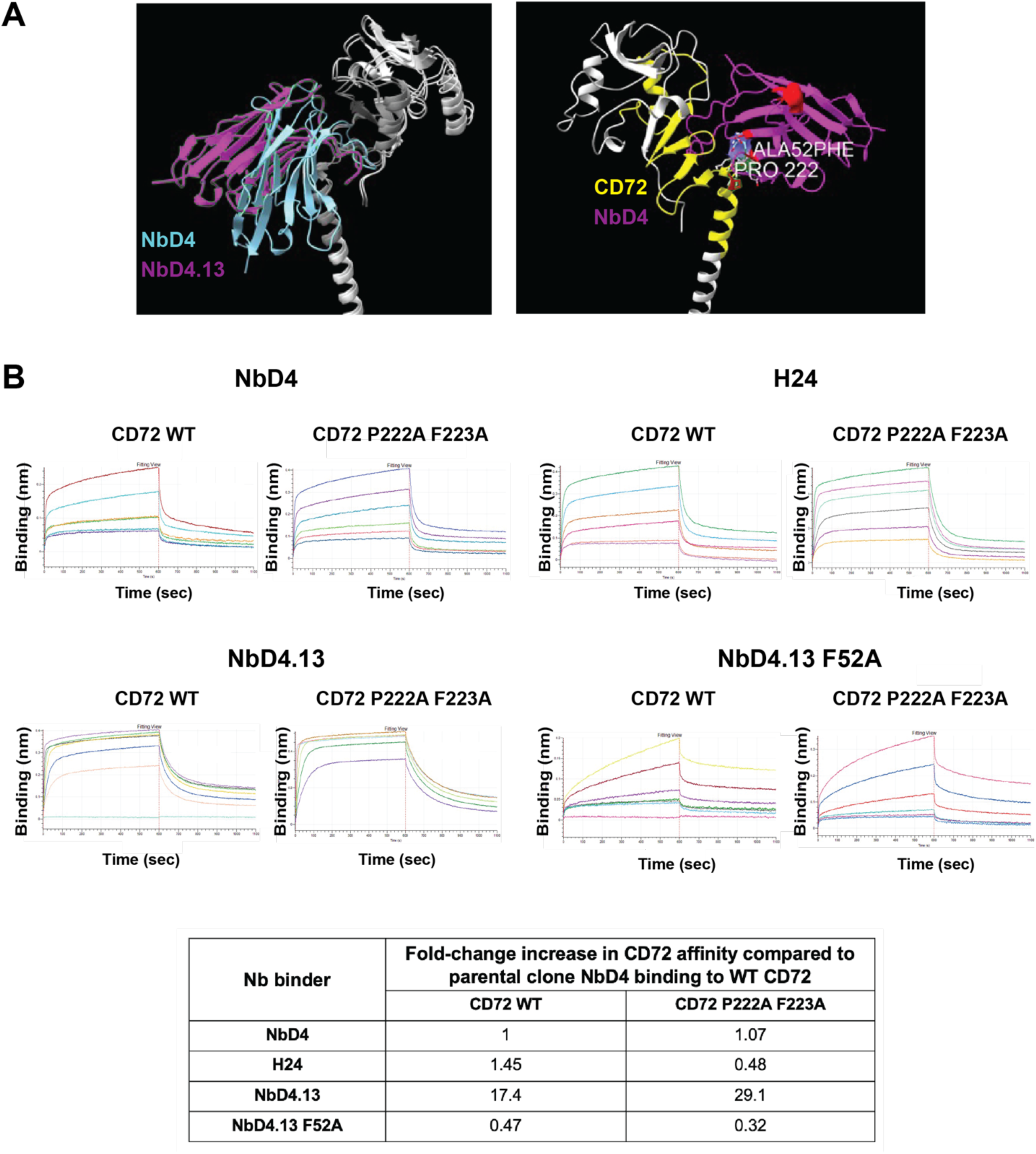
Structural modeling and biolayer interferometry studies identify critical residues in the nanobody-CD72 interaction. **A)** AlphaFold and HADDOCK modeling predicts NbD4 and NbD4.13 bind a similar epitope on the CD72 monomer: junction of CD72 coiled-coil and the lectin-like globular domain (amino acids 210-250). Only one of the three affinity maturation mutations of NbD4.13 is predicted to improve binding to CD72 (CDR2, Ala52Phe), leading to a favorable interaction of aromatic ring of NbD4.13 (Phe 52) with the proline ring (Pro 222) and adjacent aromatic phenylalanine (Phe 223) of CD72. **B)** Biolayer interferometry plots demonstrating NbD4, H24, NbD4.13 and NbD4.13 F52A nanobody binding to the biotinylated extracellular domain of CD72 (30 nM). Data are representative of a single experiment performed at multiple concentrations of input nanobody (0,0.625, 1.25, 2.5, 5, 10 µM). The table below illustrates fold-change in affinity (as measured by ratio of K_D_’s) for each combination of CD72 and tested in nanobody, all benchmarked to the parental NbD4 clone binding to CD72 WT extracellular domain. Higher values indicate increased affinity compared to NbD4 binding to CD72 WT; lower values indicate decreased affinity.

To test these structural modeling predictions, we used mutagenesis of recombinantly-expressed proteins (both recombinant nanobodies and recombinant CD72 ECD) followed by biolayer interferometry analysis. First, consistent with our modeling, we observed that reverting Phe52 on NbD4.13 to Ala significantly reduced affinity (∼37-fold) for CD72 WT ECD, and in fact created lower affinity for CD72 than the parental NbD4 clone (**Fig. 6B**). Contrary to our prediction, though, mutating Pro222 or Phe223 to Ala on CD72 ECD did not disrupt binding to NbD4.13.

Surprisingly, these mutants did decrease binding to H24 clone by ∼3.2-fold, indicating a potential difference between critical residues at the CD72 interface of H24 compared to NbD4.13. Together, these findings underscore the importance of Phe52 in CDR2 of NbD4.13 in driving increased affinity for CD72. These results also indicate potential differences in epitope binding energetics between the affinity matured NbD4.13 clone, which includes mutations in CDRs as well as the parental NbD4 nanobody framework, versus the humanized H24 clone, which only includes mutations in the parental NbD4 framework but no changes to the CDRs.

## DISCUSSION

Here we sought to investigate whether nanobody affinity maturation could improve the performance of CAR T-cells targeting CD72, with the goal of improving clinical efficacy against either CD72 low-expressing tumors, and/or to avoid tumor relapse due to CD72 downregulation. We ultimately found that affinity maturation could improve elimination of CD72 antigen-low tumors *in vitro*, but *in vivo* efficacy differences were less pronounced. We further showed that CD72 is ubiquitously expressed on B-cell cancers, and specifically found evidence that CD72 may be more likely to be preserved at the tumor cell surface after resistance to CD19-directed therapy when compared to CD22.

To our knowledge, others have not described how binder affinity maturation impacts the efficacy and properties of nanobody-based CAR T-cells. Given the commercial availability of one nanobody-based CAR T cell (ciltacabtagene autoleucel for multiple myeloma) and numerous others in preclinical or clinical development, our results here may be relevant for other emerging cellular therapies. To study the impact of low antigen density, we developed two independent isogenic models of the JeKo-1 B-NHL cell line. On the one hand, our *in vitro* results are consistent with prior studies of affinity maturation of scFv-based CAR Ts, where investigators found that increasing binding affinity could potentially improve elimination of tumors expressing low density of the target antigen. Notably, most of these studies primarily focused on *in vitro* assays^14^, but a similar dynamic was observed *in vivo* for targeting EGFR and HER2^15^. On the other hand, our *in vivo* results may be considered consistent with findings from studies with contrasting conclusions, demonstrating that very high affinity binders may actually lead to detrimental CAR functionality, whether due to impaired “serial killing”^32^, increased surface antigen trogocytosis^33^, or more rapid CAR exhaustion^34^.

Taken together, our results suggest that affinity maturation alone is not a universal solution to improving nanobody-based CAR T activity in the context of low surface antigen density. Indeed, one of the affinity matured clones we evaluated (NbD4.7) appeared to have a significant negative *in vivo* impact on anti-tumor efficacy (**Fig. 2E**). Given these hurdles, we also sought other avenues by which to enhance CD72-based tumor targeting. We found that bryostatin, a protein kinase C inhibitor, previously shown to upregulate surface CD22 on B-ALL models^16^, could also upregulate surface CD72 and synergize with CD72 nanoCARs to more efficiently eliminate tumor cells (**Fig. 5A-B**). Transcriptional profiling revealed that bryostatin treatment potently activated pathways related to B-cell receptor signaling, driving upregulation of both CD72 and CD22 (**Fig. 5D**). Bryostatin is being actively investigated in several clinical trials for treatment of Alzheimer disease and has shown a favorable efficacy versus toxicity profile^35,36^. Our findings suggest that further (pre)clinical investigation of bryostatin combination therapy with CD72 nanoCARs in B-cell cancers could be warranted.

Notably, the bryostatin results above also suggest that both CD22 and CD72 may have some common regulatory pathways. Potential common links of transcriptional regulation between *CD19*, *CD22*, and *CD72* were also suggested by recent findings of Ikaros-low mediated antigen escape after CD19 CAR T^37^. However, our flow cytometry-based findings in both cell line models and primary samples suggest that there also must be differences in regulation of surface protein expression for these B-cell markers, whether at the transcriptional or post-transcriptional level. We draw this conclusion given that CD72 appears to be more robustly maintained at the tumor cell surface after resistance to CD19-directed therapy, even in contexts where CD22 is largely lost (**Fig. 1G**). Deciphering these regulatory mechanisms will require examination in future studies.

We believe the immediate impact of this finding is to suggest that CD72 may be a more favorable second line CAR T-cell target than CD22 in the context of resistance to CD19-directed immunotherapy. For B-NHL, it is important to acknowledge that a wide variety of second-line CAR T targets after CD19 have already been described^8^. Though we found here ubiquitous CD72 expression on primary B-NHL samples, many other targets (for example, CD20) appear likely to be expressed at equal or higher surface antigen density than CD72. However, in B-ALL, the potential maintenance of surface CD72 when surface CD22 is significantly downregulated post-CD19 therapy may create real opportunities to benefit patients. This observation is particularly acute given that reported CD22-targeting CAR Ts have largely only led to short-lived remissions in refractory B-ALL^5,10^, and the leading commercial candidate CAR T targeting CD22 has recently been withdrawn from clinical consideration due to poor clinical outcomes^38^. To begin to test this hypothesis, we plan to initiate a clinical trial of our CD72 nanoCAR design in the coming months. Enrollment will be focused on B-ALL and B-NHL patients who have relapsed after prior CD19-directed therapy.

The primary limitation of our study here is that we only explored the dynamics of nanobody affinity maturation for a single target and a relatively limited set of nanobody clones. We cannot exclude the possibility that a more exhaustive investigation of different targets and/or nanobody clones could indeed identify affinity matured binders which could robustly improve both *in vitro* and *in vivo* CAR T performance in the context of low surface target expression. However, we believe that our findings here, showing that nanobody affinity maturation is not a clear-cut solution to this challenge, will remain informative for the development of future nanobody-based CAR Ts, which constitute a rapidly advancing area of investigation.

## Author Contributions

Conceptualization, data curation, formal analysis, investigation, methodology, visualization, validation, writing-original draft, writing-review and editing: A.I. Data curation, formal analysis, investigation, methodology, visualization, validation, writing-review and editing: T.R., W.C.T., B.P.E, S.W., H.G., H.T., D.G.A., A.S.K., E.R., S.Y.C., H.J., M.A.N., A.N., C.M.Y., H.W.W., S.A., S.S., R.C.L., C.C., F.S., P.P., J.A.C.S., I.T., O.Z., V.S., A.V. Resources, funding acquisition, supervision, writing-review and editing: J.M.L., M.V.M., S.P., A.V., N.N.S. Conceptualization, methodology, funding acquisition, resources, supervision, writing-original draft, writing-review and editing, guarantor: A.P.W.

## Acknowledgments

We thank patients and their families for contributing primary tissues for research purposes as used here. This work was supported by Dept. of Defense Congressionally Directed Medical Research Program Award W81XWH2210807, the UCSF Living Therapeutics Initiative, and Baszucki Lymphoma Therapeutics Initiative (all to A.P.W.), and the Multiple Myeloma Research Foundation Translational Accelerator Award (to S.P. and A.P.W.). This work was also supported by the American Society of Hematology Research Training Award for Fellows, the Chan Zuckerberg Biohub Physician-Scientist Fellowship Program, the American Society of Transplantation and Cellular Therapy New Investigator Award, the UCSF Alpha Clinic Physician Scientist Research Fellowship Award from the California Institute of Regenerative Medicine, and Alex’s Lemonade Stand Center of Excellence (all to W.C.T.), the UCSF Medical Scientist Training Program T32GM141323 (supporting A.S.K.), Harold Amos Physician-Scientist Development Program Award (to E.R.), Contrato Río Hortega by the Instituto de Salud Carlos III (CM24/00094) (to D.G-A), TERAV+ by RICORS/TERAV (RD24/0014/0003) (to A.V. and J.M-L), Intramural Research Program, Center of Cancer Research, National Cancer Institute and NIH Clinical Center, National Institutes of Health (ZIA BC 011823) (to C.M.Y. and N.N.S.). R.C.L. was supported by NIH T32AI007529 at the time of this work. Murine studies were performed at the UCSF Helen Diller Family Comprehensive Cancer Center Preclinical Therapeutic Core Facility (directed by V.S.) supported by P30CA082103. The funders of the study had no involvement in the study design, in collection, analysis and interpretation of data, writing of the report or in the decision to submit the paper for publication.

## Disclaimer

The content of this publication does not necessarily reflect the views of policies of the Department of Health and Human Services, nor does mention of trade names, commercial products, or organizations imply endorsement by the U.S. Government.

