## Supplemental methods and figures for "Affinity-matured CD72-targeting Nanobody CAR T-cells Enhance Elimination of Antigen-Low B-cell Malignancies"

Izgutdina et al.

**Supplemental Methods**

**Supplemental Table 1**

**Supplemental Figures 1-5**

**Supplementary References**

### SUPPLEMENTAL METHODS

#### Human primary samples

Human primary leukemia and lymphoma samples were obtained in adherence with local Institutional Review Board protocol, in accordance with Declaration of Helsinki.

The human primary lymphoma samples obtained from Kanazawa University with approval by its Institutional Review Board (No. 2023-283). Written informed consent was not required as consent was obtained through an opt-out process. Primary lymphoma samples were obtained from Hospital Universitario 12 de Octubre under protocol approval number 20/326. CD19 CAR T-relapsed human bone marrow/blood samples were obtained from the National Cancer Institute, treated under IRB-approved protocol (NCT03448393) and who provided consent for analysis of blood and marrow specimens for analysis of antigen expression through an IRB-approved retrospective study (NCT03827343).

#### Human cell lines and patient derived xenograft

Human cell lines JeKo-1, Toledo and Namalwa were purchased from ATCC; SC1, DOHH2 cell lines were purchased from DSMZ. Cell lines were authenticated by short tandem repeat (STR) profiling and tested negative for mycoplasma. Cells were cultured in RPMI media supplemented with 20% heat inactivated FBS and 1% penicillin+streptomycin in an incubator at $37^{o}$C and 5% $CO_2$.

Mantle cell lymphoma patient-derived xenograft (PDX) was obtained from PRoXe Dana Farber repository[1] (accession number DFBL-44685). "JeKo-1 CD72lo$^{oe}$" model was generated by CRISPR/Cas9 knock out of *CD72* gene, clonal selection of a knockout, and then re-transduction of a lentiviral construct expressing the *CD72* open reading frame. "JeKo-1 CD72 lo$^{r}$" model was generated from harvested splenocytes, after an NSG mouse implanted with WT JeKo-1 tumor relapsed after H24 CAR treatment with lower antigen density than baseline (as described in Temple et al.[2]).

#### scRNA-seq of CD19 relapsed cell lines

scRNA-seq data from B-cell malignancy cell lines (NALM-6 and REH) resistant to CD19-directed therapy was previously published[3] and re-analyzed here. Briefly, these data were generated using 10x Genomics chromium platform using 3' v2 chemistry and the raw outputs generated by running the 10x Genomics Cell Ranger software. Outputs were processed using R coding language and the Seurat Package v4.1.2. Cell line data was imported individually, and resistant and parental cell lines of the same lineage were merged into single objects (NALM-6/NALM-6-R and REH/REH-R). After standard quality control (QC) and filtering, total cells analyzed were 11,167 and 13,804 cells for NALM-6 and REH cell lines respectively. Differences in *CD72* expression between parental resistant cell lines was quantified using the FindAllMarkers function with default parameters (Wilcoxon Rank Sum test). Final UMAPS and Violin Plots showing the expression of *CD72* between different cell line conditions were generated using the FeaturePlot and VlnPlot functions in Seurat with default parameters.

**Analysis of human lymphoma sample microarray data**

Gene expression microarray data from multiple cohorts of patients with diffuse large B-cell lymphoma (DLBCL) (GSE10846 (ref.[4]); GSE23967 (ref.[5])), chronic lymphocytic leukemia (CLL) (GSE22762 (ref.[6]), and other B-non-Hodgkin lymphoma (DLBCL, follicular lymphoma (FL), Burkitt lymphoma (BL)) compared to normal B cells from non-neoplastic B lymphocytes (GSE12453 (ref.[7]) were analyzed. Expression level of a gene in a sample was determined by the average of expression values from multiple probesets on the array representing this gene. P values were calculated from two-sided Wilcoxon test. For survival analysis, the Kaplan-Meier method was used to estimate overall survival (OS) and relapse-free survival (RFS). A two-sided log-rank test was used to compare survival differences between patient groups based on whether their expression levels of a particular gene were above or below the median of the cohort. The R package 'survival' version 3.5.7 was used for the survival analysis[8].

**Molecular cloning of CAR-T vector**

DNA sequence of each CD72 CAR nanobody binder was synthesized at Twist Biosciences (South San Francisco, CA). Each binder was cloned into the lentiviral construct using NEBuilder HiFi DNA Assembly Cloning Kit (NEB, E2621S) and then expressed in competent cells NEB 5-alpha Competent *E. coli* (High Efficiency) (NEB, C2987H). DNA constructs were verified by sequencing. CAR construct DNA was amplified and isolated using QIAGEN Plasmid Plus Midi Kit (Qiagen, 12943).

**Lentiviral Vector Production**

CAR lentiviral constructs were transfected into Lenti-X 293T cells (Takara Bio, 632180) using Mirus Bio™ TransIT™-Lenti Transfection Reagent (MIR6600). Transfected Lenti-X cells were cultured for 48-72 hrs, then lentivirus was harvested and concentrated with help of Lenti-X Concentrator (Takara Bio, 631232).

**Flow cytometry assays**

At University of California, San Francisco, primary sample and cell line cell surface staining with antibodies was done per manufacturer's recommended antibody concentration, 2-5 μL in 100 μL of FACS buffer (dPBS and 2% FBS) for 15-30 minutes. Samples were washed with FACS buffer and acquired on CytoFLEX Flow Cytometer (Beckman Coulter). Compensation was done with compensation beads (UltraComp eBeadsTM Compensation Beads, Invitrogen, 01-2222-42).

For primary samples from Hospital 12 de Octubre, Spain, samples used were from hematologic neoplasms biobank registered with the code "C.0006063". Samples were stained with CD72-PE, Invitrogen (clone 3F3) and acquired in a BD FACSCanto II Flow Cytometer. For primary samples from Kanazawa University, Japan, the antibodies used for staining were CD72-PE (clone 3F3) and CD19-PE (clone HIB1) from Biolegend, the staining was done per usual protocol 5 ul/sample in FACS buffer for 15-20 min. Samples were acquired on CytoFLEX Flow Cytometer (Beckman Coulter).

For primary samples from National Cancer Institute, for flow cytometry evaluation of CD19 CAR relapsed B-ALL patient samples, specimens were received in green-top sodium heparin collection tubes. Specimens were processed within 12 hours of collection. Whole blood lysis was performed using ammonium chloride. The sample was then stained with a panel of antibody cocktails for 30 min at room temperature. Samples were acquired using an 8-color multiparametric approach on a 3-laser FACS Canto II (BD Biosciences, San Jose, CA) with

DiVA software.  Flow cytometry data was analyzed with FCS Express software (DeNovo Software, Los Angeles, CA).

**Exhaustion and memory staining**

Exhaustion and memory analysis was performed by staining CAR T-cells with CD3, PD1, LAG3, TIM3 antibodies after 24 hour coculture with JeKo-1 tumor cells at 1:1 E:T ratios. Memory staining was done with CD62L, CD45-RA antibodies after 24 hour coculture with JeKo-1 tumor at 1:1 Effector:Tumor (E:T) ratio. Samples were acquired on CytoFLEX Flow Cytometer (Beckman Coulter). Please See Supplementary Table 1 for information on antibody clones and fluorophores.

**Activation and degranulation assay**

Activation and degranulation staining was performed after 6 hour coculture at 1:1 E:T ratio in the presence of 2 uL of anti-CD107a antibody and 1X Monensin (eBioscience, 420701). Cells were washed and subsequently stained with CD69 antibody. Percent positive cells for CD107a and CD69 were assessed as marker of degranulation and activation, respectively. Samples were acquired on CytoFLEX Flow Cytometer (Beckman Coulter).  See Supplementary Table 1 for information on antibody clones and fluorophores.

***In vitro* cytotoxicity assays**

Untransduced T cells or CAR T-cells were incubated with JeKo-1, Toledo, Namalwa, SC-1 or Jeko-1 CD72lo$^{oe}$ tumor cell lines to measure cytotoxicity. All tumor cell lines were engineered to stably express luciferase. For each assay, 0.1e6 tumor cells were seeded into 96 well plates. CAR T-cells were added at 1:1, 1:3 and 1:10 E:T ratios in total volume of 200 μL complete RPMI media in triplicate. Cells were incubated for 24 hours at 37 $^{o}$C. The next day 2 μL of D-Luciferin solution (Gold Biotechnology, LUCK-1G) was added to each well to reach 15 μg/mL concentration. The plate was incubated for 10 minutes and bioluminescence signal was assessed using GloMax Explorer (Promega). Bioluminescence of each well was normalized to the average bioluminescence of untransduced T cell and tumor coculture at indicated E:T ratio to derive percent cytotoxicity.

**Incucyte assay**

Untransduced T cells or CAR-T cells were cultured in 96 well plates with JeKo-1 or JeKo-1 $CD72lo^{oe}$ cell lines engineered to express mCherry at 1:1, 1:3 and 1:10 E:T ratios in a total volume of 200 μL complete RPMI media. The co-culture plate was imaged every 4 hours in Incucyte Live cell analyzer (Sartorius) for a total of 4-6 days. Incucyte Live-Cell Analysis software (Sartorius) was used for data analysis. Data was normalized to initial reading of respective well to calculate cytotoxicity index at each time point and plotted over time. In the experiment with JeKo-1 $CD72lo^{r}$ cell line, raw values of each well reading was plotted to represent tumor cells at 36 hour coculture time point.

**Murine studies**

NSG (NOD.Cg- $Prkdc^{scid}$ $Il2rg^{tm1Wjl}$/SzJ, Jackson Laboratories) mice were used for all murine studies. For JeKo-1 WT studies, mice were implanted via tail vein injection 1e6 tumor cells stably expressing luciferase. To assess tumor burden, mice were injected with D-luciferin and imaged with IVIS imaging system (Perkin Elmer In Vivo Imaging System, Caliper Life Sciences) to quantify bioluminescence signal. Prior to initiating treatment, mice were randomized to study arms to ensure approximately equal tumor burden across arms. 3.5e6 Empty CAR, CD19 or CD72 CAR T-cells were injected 7 days after tumor implantation.

For initial PDX experiment, mice were implanted with 1e6 DFBL-44685 mantle cell lymphoma PDX cells intravenously and injected with 5e6 Empty, H24, NbD4.7 or NbD4.13 CAR T-cells 10 days after tumor injection. Mice treated with empty CAR were sacrificed at humane end point after injection. For second PDX experiment at higher tumor burden, mice were injected with 2e6 cells and followed by untransduced T-cells ($n$=7), H24 or NbD4.13 CAR T-cell ($n$=4 each) injection at 2 weeks. The mice were monitored with blood tumor and CAR measurement at 5 weeks and spleen ultrasound at 6 weeks after tumor injection. For JeKo-1 $CD72lo^{oe}$ mouse experiment, 0.5e6 tumor cells were injected followed by 5e6 Empty, CD19, H24, NbD4.13 or NbD4.13-H24 CAR T-cells one week later. Humane end point necessitating sacrifice was used to determine survival in all murine studies.

**CAR T and tumor measurement in murine peripheral blood**

Murine blood samples were collected every 2 weeks after CAR injection. Mouse blood was treated with 1X RBC lysis buffer (Biolegend, 420301) for 10 minutes, subsequently the cells were washed with dPBS and stained with anti-CD3 or anti-CD45 and anti-CD20 antibodies (Supplementary Table 1). Cells were acquired on CytoFLEX Flow Cytometer (Beckman Coulter). Flow cytometry data was analyzed on FlowJo software (version 10.10.0) with gating on CD3+ and GFP+ cells or CD45+ GFP+ cells to assess for CAR+ T cell population percentage. Cells were gated on CD45+ and CD20+ for lymphoma tumor cell assessment.

**Repetitive stimulation assay**

Untranduced T cells or CAR-Ts were cultured with JeKo-1 or JeKo-1-CD72lo$^{oe}$ cell lines at 1:1 ratio. 1e6 of the cell coculture was transferred to a new plate with respective cell line at 1:1 ratio every 3 days. Cells were used for Incucyte analysis at 2$^{nd}$ and 5$^{th}$ stimulation with tumor.

***CD72* Knockout by CRISPR-Cas9**

sgRNA guides (Synthego Corporation) and Cas9 protein (Macrolab, University of California, Berkeley) were incubated at 2:1 ratio at 37C for 15 minutes. 1e6 JeKo-1 cells were resuspended in SF nucleofector and supplement solution (SF Cell Line 4D-NucleofectorTM X Kit S, Lonza, V4XC-2032) with the addition of Cas9/sgRNA mixture.

The cell line was nucleofected on 4D-Nucleofector X Unit (Lonza, AAF-1003X) using DJ-105 nucleofection program. Cells were then recovered in 80 uL of RPMI media in 37C incubator for 15 minutes and then transferred to a tissue culture plate. Successful knockout was confirmed with flow cytometry staining for CD72 surface expression (FACSARIA III flow cytometer (BD Biosciences)). The knockout cells were initially sorted by FACS to isolate a CD72 negative population. A subsequent round of sorting was done for clonal isolation of CD72 knockout (KO) clones. CD72 targeting guide sgRNA was obtained from Brunello library[9]: *GAGTCAGGAAGCACTACAGG*.

**Cytokine secretion assay**

CAR-Ts were cultured with JeKo-1 or JeKo-1 CD72-low tumor cell lines in complete RPMI media at 1:1 E:T ratio for 24 hours. Cell supernatant was harvested in triplicates for each CAR condition. Supernatant was diluted 1:1 in RPMI media, snap frozen in liquid nitrogen and stored at -80 $^{o}$C. Cytokine samples were analyzed by Eve Technologies (Calgary, Alberta) using Luminex xMAP technology for multiplexed quantification of 14 Human cytokines. Luminex™ 200 system (Luminex, Austin, TX, USA) by Eve Technologies Corp. was used for multiplexing analysis. Cytokine markers were measured using Eve Technologies' Human High Sensitivity 14-Plex Discovery Assay® (MilliporeSigma, Burlington, Massachusetts, USA). The 14-plex consisted of GM-CSF, IFNγ, IL-1β, IL-2, IL-4, IL-5, IL-6, IL-8, IL-10, IL-12p70, IL-13, IL-17A, IL-23, TNF-α, anaylzed in triplicate. Assay sensitivities of these markers range from 0.11 – 3.25 pg/mL for the 14-plex. Individual analyte sensitivity values are available in the MilliporeSigma MILLIPLEX® MAP protocol. Point-to-Point Semi Log regression was applied to estimates of IFNγ analytes. This type of regression enables extrapolation and provides number values to all out of range data.

**CAR T avidity study**

CAR T-cell avidity was assessed with z-Movi cell Avidity Analyzer v17 (LUMICKS). The chip was activated with NaOH and coated with poly-L-lysine to allow target cell adhesion. The chip was then rehydrated with serum free RPMI media and stored at 37$^{o}$C. Jeko-1 or Jeko-1 CD72lo$^{oe}$ cells were then evenly seeded at 130e6/mL on the chip and were incubated for 1 hour. Media was refreshed and cells were incubated for 1 additional hour to ensure strong chip adhesion.

T cells were stained with CellTrace Far Red kit (Thermo Fisher, C34564) to differentiate from target cells and untransduced T cells and Empty CAR were used as negative controls. The order of CAR Ts assayed was randomized between chips. CAR-Ts were incubated with tumor for 5 minutes to allow interaction and then a ramped acoustic force from 0-1000 pN was applied vertically to the cells over the course of 3 minutes. The cells were then imaged using the z-Movi cell Avidity Analyzer and analysis of the binding force was performed using the Oceon™ software (LUMICKS). All experiments were performed in duplicate or triplicate.

**Bryostatin and Azacytidine Treatment of Cell Lines**

Bryostatin (Bryostatin-1, Sigma, 203811)was reconstituted in DMSO to 1 μM stock solution. SC-1, DOHH-2 and JeKo-1 cell lines were cultured in the presence of varying concentrations of bryostatin (0.1 nM and 1 nM) or equal volume of DMSO in complete RPMI media for 3 days. On Day 3 cells were stained with CD72, CD19 and CD22 antibodies (see Supplementary Table 1 for antibody clones) and BD Horizon™ Fixable Viability Stain 450 and analyzed on a BC Cytoflex flow cytometer. SC-1 and DOHH-2 cell lines were cultured in the presence of azacytidine (5-Azacytidine, Selleckchem, S1782) at 1.2, 2.5, 5, 10 μM concentration or equal volume of 0.1% DMSO. Cells were first treated every day for three days at indicated concentrations of drug or 0.1% DMSO and then cultured for four days in complete RPMI media without drug. On day 8, cells were stained with CD72, CD19 and CD22 antibodies and analyzed by flow cytometry.

**Bulk RNA sequencing of cell lines**

SC-1 cells were cultured in the presence of DMSO or 1 nM bryostatin for 72 hrs with fresh media with drug was added every 24 hours. Qiagen RNeasy Mini kit was used for RNA extraction from SC-1 cells. Sample QC was performed with Tapestation (Agilent) to identify samples with RIN>7. 200 ng qualified RNA from each sample was processed for library preparation. In brief, mRNA enrichment was performed on total RNA using oligo(dT)-attached magnetic beads. The enriched mRNA with poly(A) tails was fragmented using a fragmentation buffer, followed by reverse transcription using random N6 primers to synthesize cDNA double strands. The synthesized double stranded DNA was then end-repaired and 5'-phosphorylated, with a protruding 'A' at the 3' end forming a blunt end, followed by ligation of a bubble-shaped adapter with a protruding 'T' at the 3' end. The ligation products were PCR amplified using specific primers. The PCR products were denatured to single strands, and then single-stranded circular DNA libraries were generated using a bridged primer. The constructed libraries were quality-checked and sequenced after passing the quality control. The library was amplified with phi29 to make DNA nanoball (DNB) which had more than 300 copies of one molecule. The DNBs were load into the patterned nanoarray and pair end 150 bases reads were generated in the way of sequenced by synthesis. Sequencing was performed using the DNBSEQ-T7 (MGI Tech) platform. Raw sequencing reads were subjected to quality control (QC) to determine whether the sequencing data were suitable for subsequent analysis. After QC, the clean reads were aligned to the reference sequences.

**Analysis of bulk RNA sequencing of cell lines**

The initial RNA-seq data was processed using SOAPnuke (version 1.5.6 (ref.[10])) with the parameters: (-l 15 -q 0.2 -n 0.001 -G) to remove low-quality reads, adapter sequences, and PCR duplicates. The filtered reads were aligned to the reference human genome (GRCh38.p13) using a spliced aligner Hisat2 (version 2.2.1 (ref.[11]) with the parameters: --sensitive --no-discordant --no-mixed -I 1 -X 1000 -p 8 --rna-strandness RF. Gene-level expression values, including raw read counts, FPKM (fragments per kilobase of transcript per million fragments mapped), and TPM (transcripts per million reads), were quantified using RSEM (version 1.2.28 (ref.[12]) with the parameters: (-p 8 --forward-prob 0). The raw read counts were normalized and differential expression between two groups (Bryostatin treated, n=3; or DMSO treated, n=3) were calculated using DESeq2 (version 1.40.2 (ref.[13])). UMAP dimensionality reduction was performed on gene expression FPKM and TPM values using all genes. The significantly differentially expressed genes (q-value<0.05) were presented by heatmaps and volcano plots. The R software[8] was used with the umap and pheatmap libraries. The pathway analysis on the differentially expressed genes was conducted using DAVID software[14]. Raw sequence files are deposited at the Gene Expression Omnibus repository (see Data Availability).

**scRNAseq of mouse tumors**

Human mantle cell lymphoma PDX tumor (DFBL-44685) was implanted at 1e6 cells/mouse, mice were then treated with Empty, H24 or NbD4.13 CAR at 5e6 CAR+ cells per mouse on day 10. Spleens were collected at day 70 after tumor implantation. Tumor cells were isolated from mouse spleens and sorted with gating on human CD45+ positive cells. The samples were processed using 10X chromium platform (Chromium Next GEM Immune Profiling 5'HT kit v2, 1000374) according to protocol. Illumina sequencing was performed on the mouse spleen samples. The reads were aligned and counted using STARSolo version 2.7.11b, with --featureType Gene, and --soloCellFilter EmptyDrops_CR. The genome and GTF were taken from the cellranger ref-GRCH38-2024-A, which is derived from GENCODEv44. To generate feature-barcode matrices analysis was conducted in R (v4.1.2) with Seurat (v4.3.0)[15]. Quality control excluded cells expressing fewer than 500 genes, fewer than 1,000 total RNA counts, more than 50,000 features, or with >10% mitochondrial content. Data were normalized, and highly variable features were identified for downstream analysis. Batch effects were corrected with the Harmony[16] package in RStudio (v0.1.1), grouping by experimental condition, and embeddings were used for UMAP visualization and clustering (resolution = 0.5). Gene symbols were annotated using biomaRt. Differentially expressed genes (DEGs) between clusters were identified with a log fold change threshold >0.25 and filtered for average log fold change >0.5. Visualization was performed with ggplot2[17] (v3.5.1) and gridExtra[18] (v2.3). Analysis code is available in the GitHub repository at [https://github.com/BonellPatinoE/CD72_scRNAseq-in-mantle-cell-lymphoma.git].

***TRAC* knockout for *in vivo* studies**

To avoid development of graft-vs-host-disease, CAR T-cells in murine studies versus human PDX DFBL-44685 and JeKo-1 CD72lo$^{oe}$ cell line underwent CRISPR/Cas9 knock out of T-cell receptor alpha chain (*TRAC*) exon 1 (sgRNA sequence: CAGGGTTCTGGATATCTGT), as previously described by others[19]. Nucleofection with Cas9 and sgRNA was done on day 1 after thaw using P3 Primary Cell 4D X Kit (Lonza, V4XP-3032) and 4D-Nucleofector X Unit (Lonza, AAF-1003X), using EO-115 nucleofection program. After electroporation, CAR T-cells were enriched for successful *TRAC*-KO with human CD3 MicroBeads kit (Miltenyi Biotec, 130-050-101).

**Nanobody epitope modeling**

Nanobody structures were generated in IgFold (COSMIC2 server[20]) using standard parameters[21]. The structure for human CD72 was generated in AlphaFold2. The relaxed CD72 structure was docked in HADDOCK Version 2.4 (ref.[22,23]) following the basic antibody-antigen docking guide using nanobody residues 1-122 and the extracellular domain of CD72 (aa 220-359) as active residues. The number of structures for semi-flexible and final refinement were increased to 400. The best scoring models (lowest/most stable HADDOCK score) were visualized in UCSF ChimeraX and potentially interacting residues were visualized.

**Molecular cloning and DNA vector construction for recombinantly-expressed proteins**

The expression plasmid encoding the anti-CD72 nanobody sequence was synthesized by Twist Biosciences. The construct included an N-terminal PelB signal sequence for periplasmic expression in *E. coli* BL21(DE3) cells and was cloned into the pET29b vector with a C-terminal hexahistidine tag (6xHis) in *NdeI* and *XhoI* restriction sites. DNA encoding the extracellular domain of CD72 (amino acids 117–359) or a mutant variant (P222A F223A) with upstream IL-2 secretion signal with an N-terminal AviTag for site-specific biotinylation and a human IgG CH2–CH3 constant region (Fc) was synthesized by Twist Bioscience. The DNA fragments were cloned into a pcDNA-based mammalian expression vector using Gibson Assembly (NEB, E2611S) at *XbaI* and *AgeI* restriction sites. The recombinant plasmid sequence was verified by DNA sequencing. The final construct was transformed into *E. coli* NEB 5-alpha competent cells and plasmid DNA was isolated using the QIAGEN Plasmid Plus Midi Kit (12943) for transfection into Expi293 cells.

**Transfection, protein expression, and protein purification**

Expi293 cells were transfected at 3 million cells/mL density using the ExpiFectamine™ 293 Transfection Kit (Thermo Fisher Scientific, A14524) following the manufacturer's instructions. Five days post-transfection, the culture supernatant was harvested by centrifugation at 4000 rpm for 10 minutes and clarified by filtration through a 0.45 μm membrane. The pH was adjusted to 7.4 using PBS. The clarified supernatant was loaded onto a HiTrap Protein A affinity column (Cytiva, 17040201) and pre-equilibrated with PBS (pH 7.4). After washing the column with 5-column volume PBS, the bound protein was eluted with 0.1 M acetic acid. The eluate was immediately neutralized with 1M Tris buffer (pH 11) and subjected to buffer exchange into PBS (pH 7.4) using 10 kDa MWCO centrifugal filters (Amicon Ultra, UFC9010). Purified proteins were aliquoted and flash-frozen for storage at −80 °C for long-term use. Protein samples were prepared in 2X SDS loading dye with  DTT and analyzed by SDS-PAGE. Protein concentration was determined using absorbance at 280 nm (A280) on a NanoDrop spectrophotometer (Thermo).

**His-tagged nanobody expression and purification:**

For each purification, *E. coli* BL21 (DE3) were grown in Terrific Broth (Research Products International) and induced after reaching $OD_{600}$ 0.6–0.8 with 1 mM IPTG (Sigma, I6758) at 24 °C for 16 hours. The induced cells were harvested by centrifugation and resuspended in 50 mL of lysis buffer containing 0.5 M sucrose, 200 mM Tris-HCl (pH 8.0), 0.5 mM EDTA, protease inhibitor cocktail (Sigma, 11836170001), lysozyme (1 mg/mL), and DNase I (10 μg/mL) and stirred at room temperature for 30 minutes. Periplasmic extraction was performed by osmotic shock, adding 100 mL of deionized water and stirring for 45 minutes at RT. The lysate was adjusted to a final concentration of 150 mM NaCl, 2 mM $MgCl_2$, and 20 mM imidazole and then centrifuged at 20,000g for 20 min at 4 °C. The clarified supernatant was applied to a gravity column containing 3 mL of Ni Sepharose 6 Fast Flow resin (GE Healthcare, 17531802). The resin was washed with a high-salt buffer (20 mM HEPES, pH 7.5, 500 mM NaCl, 20 mM imidazole), followed by a second wash with low-salt buffer (20 mM HEPES, pH 7.4, 100 mM NaCl, 20 mM imidazole). The nanobody protein was eluted using 20 mM HEPES (pH 7.4), 100 mM NaCl, and 250 mM imidazole. The eluted protein was buffer-exchanged into PBS (pH 7.4) using 10 kDa MWCO centrifugal filters (Amicon Ultra, UFC9010). As mentioned, protein samples were prepared in 2× SDS loading dye containing DTT and analyzed by SDS-PAGE. Purified nanobodies were stored at 4 °C for long-term use.

**Biolayer interferometry (BLI) for protein-protein interaction analysis**

Biolayer interferometry (BLI) experiments were conducted using an Octet RED384 instrument (ForteBio, Sartorius). Biotinylated CD72 extracellular domain (ECD) or its mutant variant (P222A F223A) was immobilized on streptavidin (SA) biosensors (ForteBio) by dipping the sensors into a solution of the biotinylated proteins until a binding signal of approximately 0.6 nm was achieved. To prevent non-specific binding, the biosensors were subsequently blocked with 10 µM free biotin. Purified nanobody proteins were used as analytes and prepared in the buffer of PBS (pH 7.4) with 0.05% Tween-20 and 0.2% BSA (PBSTB). A range of increasing nanobody concentrations was titrated to assess binding kinetics. Association and dissociation steps were monitored by sequentially dipping the biosensors into the analyte solution, followed by buffer-only wells. Data was processed and analyzed using ForteBio Octet analysis software and the dissociation constant ($K_D$) was determined by fitting the data to a 1:1 Langmuir binding model suitable for monovalent interactions.

**Data and Code Availability**

scRNAseq analysis code is available in repository at https://github.com/BonellPatinoE/CD72_scRNAseq-in-mantle-cell-lymphoma.git.

scRNAseq data generated here is deposited at Gene Expression Omnibus (GEO):

https://www.ncbi.nlm.nih.gov/geo/query/acc.cgi?acc=GSE290685

Accession: GSE290685. Reviewer token: abctcuquzlqpzad

RNA-seq data generated here is deposited at Gene Expression Omnibus (GEO):

https://www.ncbi.nlm.nih.gov/geo/query/acc.cgi?acc=GSE290380.

Accession: GSE290380. Reviewer token: ubkfsygqfvevtml.

### SUPPLEMENTARY FIGURES

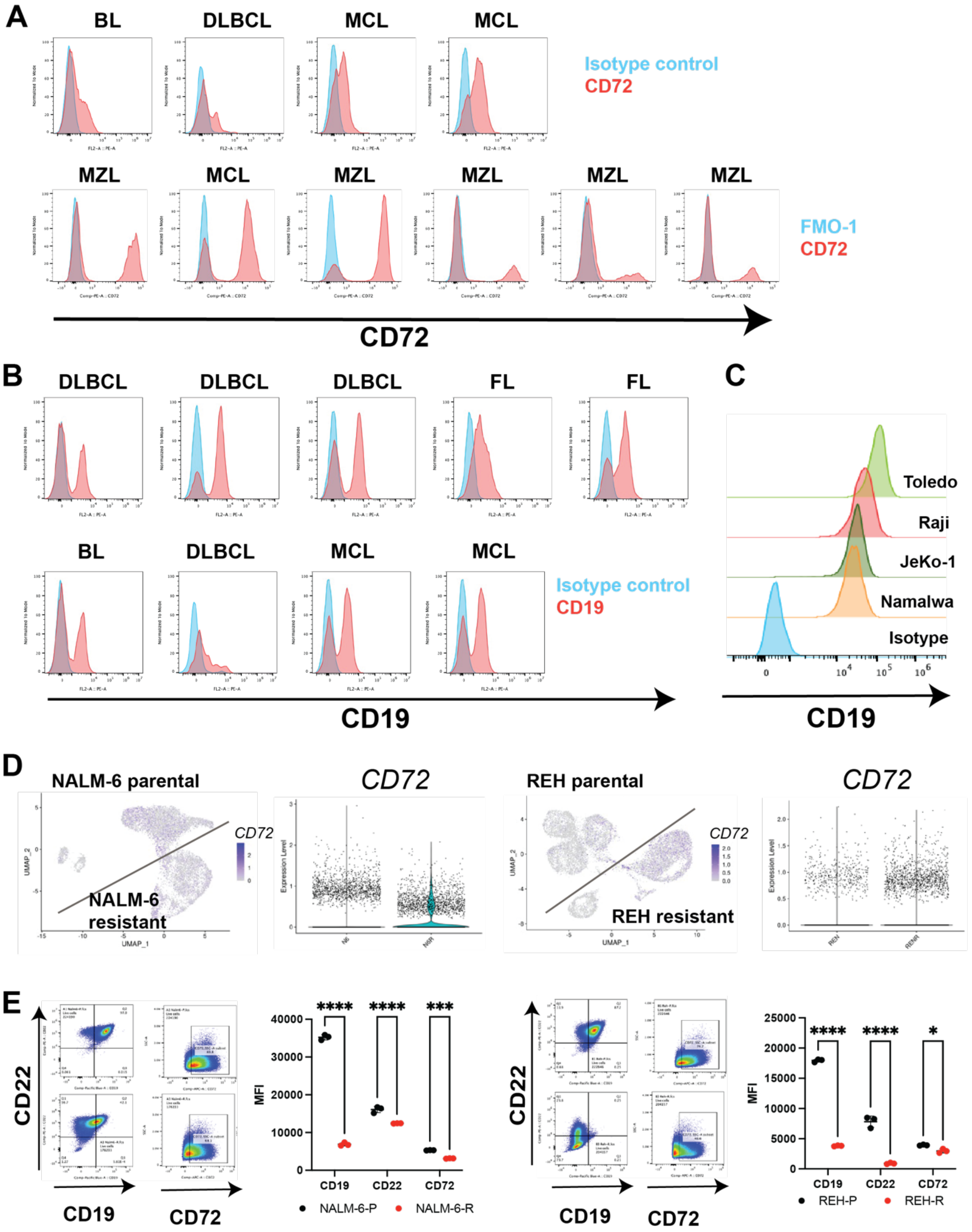


**Supplementary Figure 1: CD72 is a target present on primary lymphoma samples and retained post-CD19 therapy relapse. A)** Flow cytometry of primary lymphoma tumor samples with measurement of CD72 expression. For lymphoma samples from Kanazawa University,

Japan, (top panel), isotype control was used as negative control and for samples from Hospital Universitario 12 de Octubre, Madrid, Spain (bottom panel), fluorescence minus one was used as negative control . **B)** Flow cytometry of primary lymphoma tumor samples from Kanazawa University, Japan with measurement of CD19 expression. Compared to isotype control. **C)** Flow cytometry of lymphoma cell lines with measurement of CD19 expression. Compared to isotype control. **D)** UMAP depicting *CD72* expression in wild type and CD19-directed therapy resistant NALM-6 and REH cell lines (resistant cells developed and scRNA-seq data obtained as described in ref.[3]). Violin plots comparing *CD72* expression between WT and CD19-resistant NALM-6 and REH cells. **E)** Flow cytometry plots and MFI data of WT and CD19 resistant NALM-6 and REH cell lines depicting expression of CD19, CD22 and CD72 before and after developing resistance.

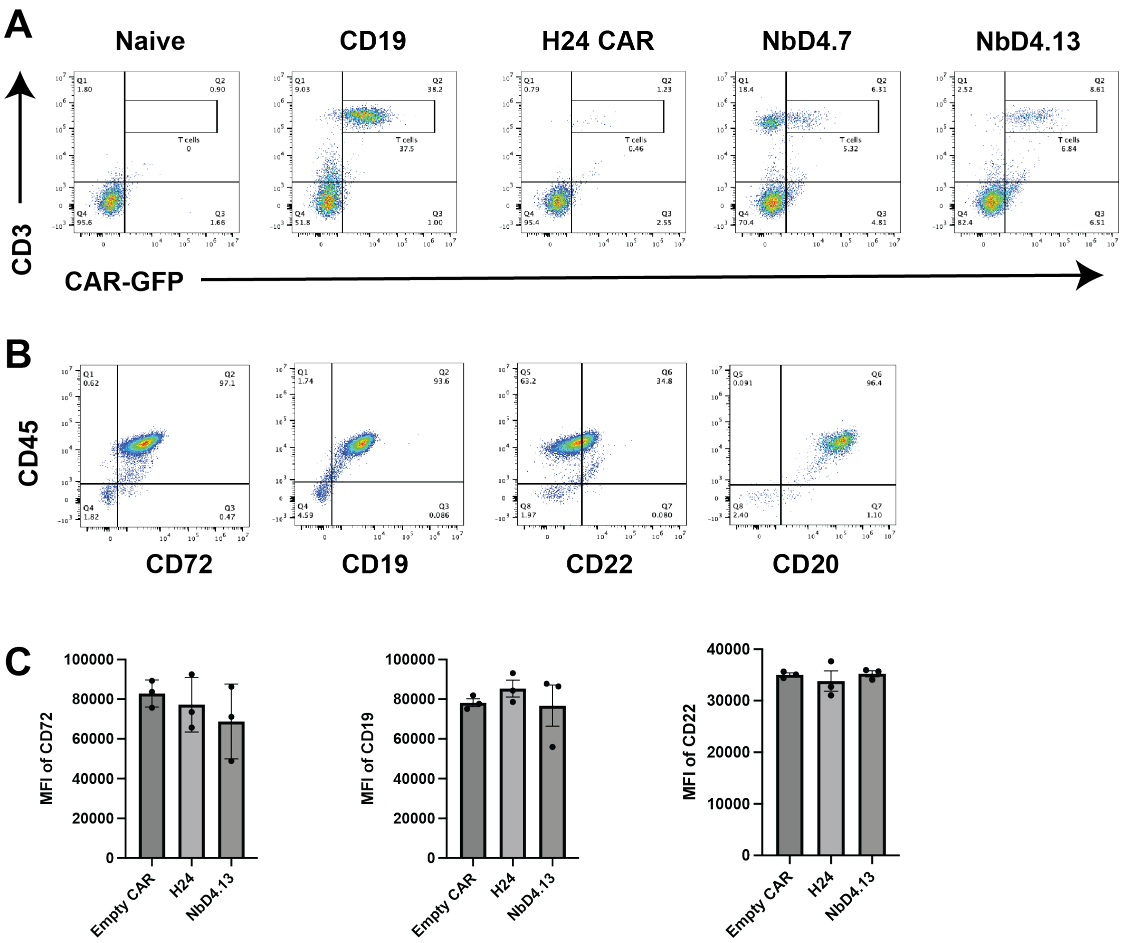


**D**

| | CDR1 | | CDR2 | | CDR3 | |
|---|---|---|---|---|---|---|
| **NbD4** | QVQLQESGGGLVQAGGSLRLSCAASG**TISPIDI** | MGWYRQAPGKERE**FVAAIALGGN** | | TYYADSVKGRFTISRDNAKNTVYLQMNSLKPEDTAVYYCA**VGYVDKWDDSNYHT** | | YWGQGTQVTVSS |

```
                        CDR1             CDR2                                          CDR3
NbD4         QVQLQESGGGLVQAGGSLRLSCAASGTISPIDIMGWYRQAPGKEREFVAAIALGGNTYYADSVKGRFTISRDNAKNTVYLQMNSLKPEDTAVYYCAVGYVDKWDDSNYHTYWGQGTQVTVSS
H24          EVQLVESGGGLVQPGGSLRLSCAASGTISPIDIMSWYRQAPGKEREFVAAIALGGNTYYADSVKGRFTISRDNSKNTLYLQMNSLRAEDTAVYYCAVGYVDKWDDSNYHTYWGQGTQVTVSS
NbD4.13      QVQLVESGGGLVQPGGSLRLSCAASGTISPIDQMGWYRQAPGKEREFVAAIFLGGNTYYADSVKGRFTISRDNAQNTVYLQMNSLKAEDTAVYYCAVGYVDKWDDSNYHTYWGQGTQVTVSS
NbD4.13-H24  EVQLVESGGGLVQPGGSLRLSCAASGTISPIDQMSWYRQAPGKEREFVAAIFLGGNTYYADSVKGRFTISRDNSQNTLYLQMNSLRAEDTAVYYCAVGYVDKWDDSNYHTYWGQGTQVTVSS
```

**Supplementary Figure 2. CAR-T peripheral blood and PDX tumor model flow cytometry.** **A)** Mouse peripheral blood flow cytometry staining for hCD3+/GFP+ CAR T-cells in an *in vivo* study versus JeKo-1 WT tumor cells (corresponding to study shown in **Fig. 2E** of main text). **B)** Flow cytometry profiling of CD20, CD19, CD72, CD22 tumor markers on MCL PDX DFBL-44685 at untreated baseline (corresponding to study shown in **Fig. 2H** of main text). **C)** Flow cytometry data demonstrating largely equivalent CD72, CD19 and CD22 surface antigen expression both before and after CAR treatment and relapse in PDX DFBL-44685 bearing mice. **D)** Amino acid sequence of CD72 CAR-Ts: parental NbD4, humanized H24, affinity matured NbD4.13 and affinity matured + humanized binder NbD4.13-H24.

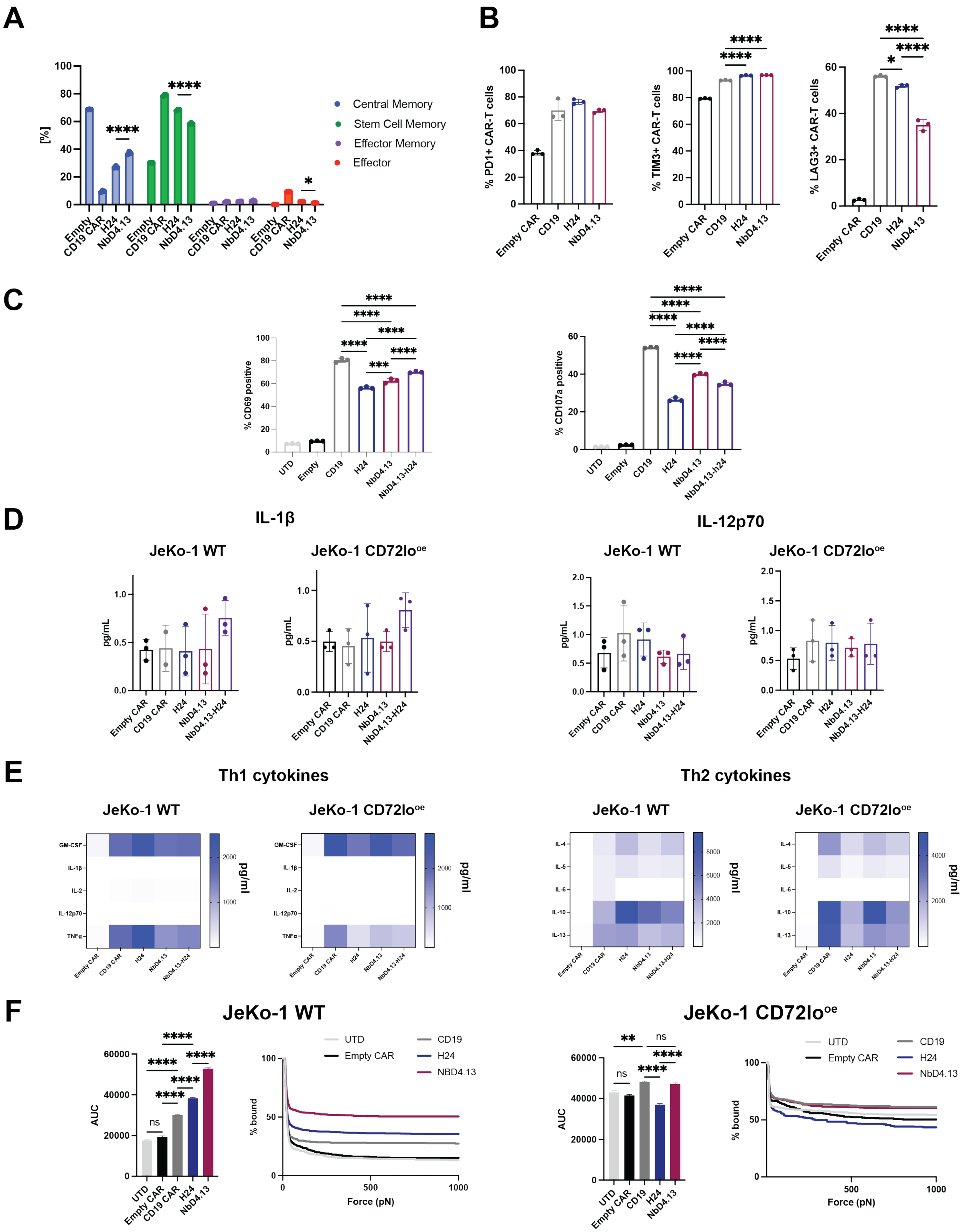


**Supplementary Figure 3. In vitro characterization of affinity matured CD72 CAR-T cells.**

**A)** Memory phenotype of CD72 CAR-Ts assessed by flow cytometry staining with CD45RA and CD62L after 24hr coculture with JeKo-1 WT tumor at 1:1 effector:tumor (E:T) ratio. The bar graph shows percentages of central memory (CD45RA- CD62L+), stem cell memory (CD45RA+CD62L+), effector memory (CD45RA- CD62L-) and effector (CD45RA+ CD62L-) cell phenotypes. *p*-values obtained by 2-way ANOVA with Uncorrected Fisher's LSD (ns $p>0.05$, *$p<0.05$, **$p<0.01$, ***$p \leq 0.001$, ****$p \leq 0.0001$). **B)** Exhaustion phenotype (PD-1, TIM-3, LAG-3 expression) of CD72 CAR-Ts assessed by flow cytometry after 24hr coculture with JeKo-1 WT tumor at 1:1 effector:tumor (E:T) ratio. **C)** Activation (CD69) and degranulation (CD107a) markers assessed by flow cytometry after 6 hr stimulation with JeKo-1 WT tumor at 1:1 effector:tumor (E:T) ratio. **D)** Supernatant cytokine profiling of CD72 CAR-Ts after 24hr exposure to Jeko1 WT and Jeko1-low tumor models at 1:1 effector:tumor (E:T) ratio, with focus on IL-1β and IL-12, showing no significant differences. Analysis in B-E performed by one-way ANOVA with Tukey's multiple comparisons test (ns $p>0.05$, *$p<0.05$, **$p<0.01$, ***$p \leq 0.001$, ****$p \leq 0.0001$). **E)** Heat map of broader analysis of T-helper 1 (Th1) and T-helper 2 (Th2) multiplexed cytokines (from same assay in E)). $n=2$ or 3 replicates. **F)** Acoustic force microscopy assessment of binding avidity of CD72 CAR-Ts to JeKo-1 WT and JeKo-1 CD72lo$^{oe}$ cell lines. Area under the curve (AUC) and percent bound cells are plotted. Data represents mean with n=2 or 3, in a single experiment.

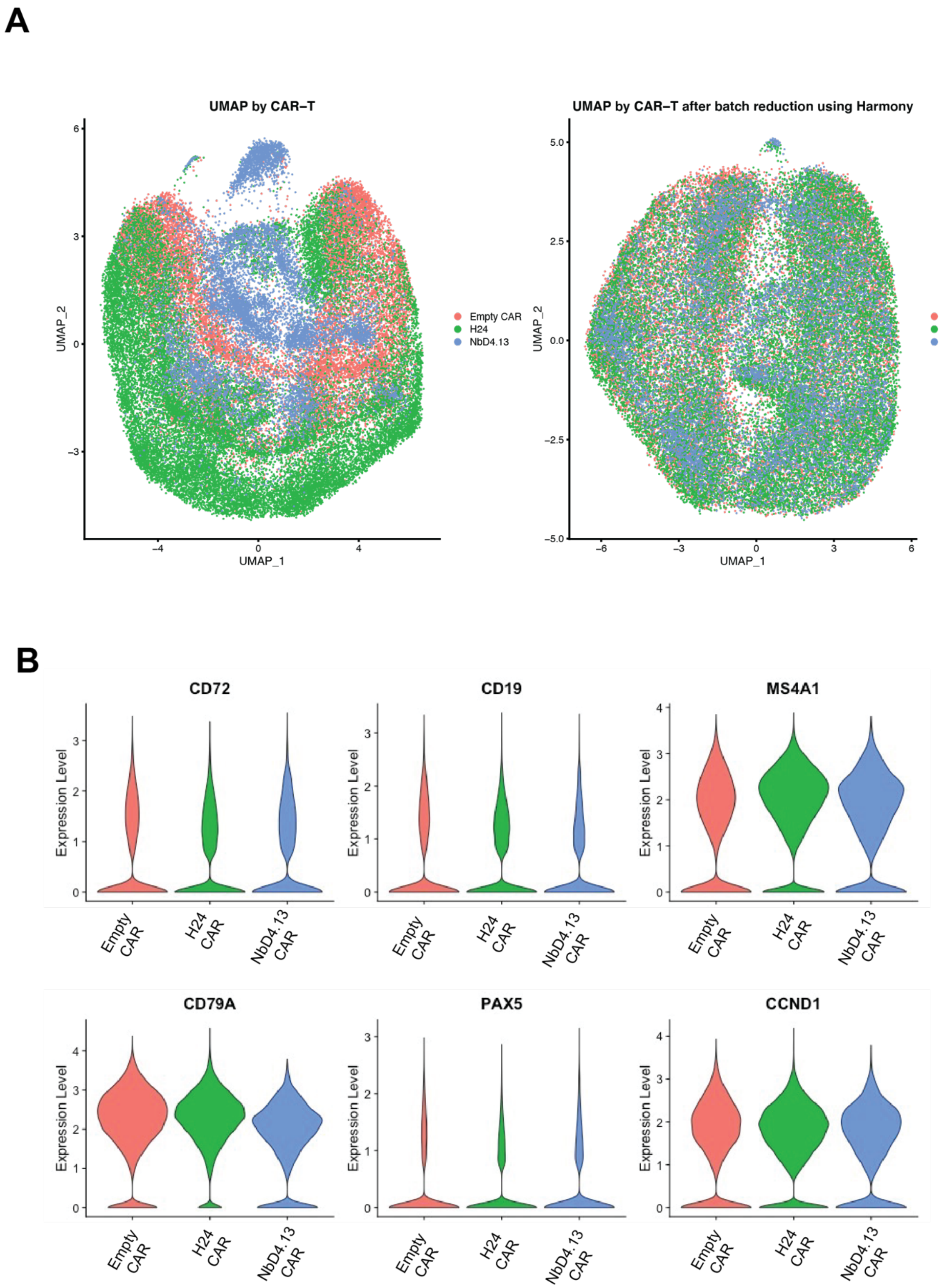


**Supplementary Figure 4. scRNA-Seq of humanized and affinity matured CD72 CAR relapsed mouse spleens demonstrates similar tumor expression patterns after relapse. A)**

UMAP plots show gene expression in CAR-T groups before and after Harmony[16] integration to resolve batch effects. **B)** Violin plots depicting canonical markers of mantle cell lymphoma expressed in Empty, H24 and NbD4.13 treated groups.

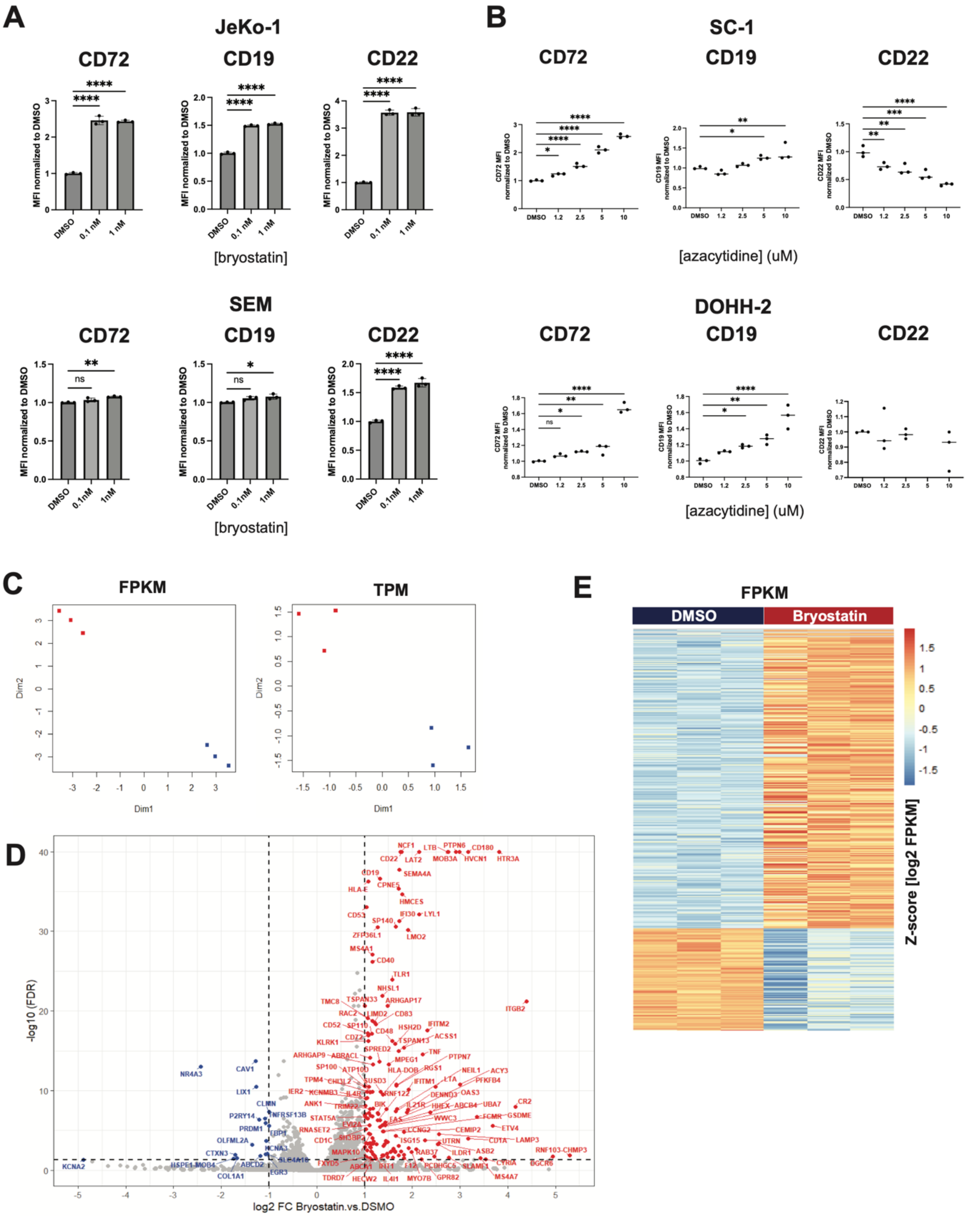

**Supplementary Figure 5. Bryostatin upregulates targets by generalized upregulation of B-cell specific signatures. A)** JeKo-1 (top panel) and SEM (bottom panel) cells treated with 0.1%

DMSO or increasing concentrations of bryostatin for 72 hours. Surface antigen density of CD72, CD19, CD22 profiled by flow cytometry and normalized to DMSO-treated cells. **B)** SC-1 (top panel) and DOHH-2 (bottom panel) cell lines treated with 0.1% DMSO or indicated concentrations of azacitidine for 4 days. Flow cytometry profiling of target surface expression after treatment with drug is normalized to DMSO-treated cells. *p*-value analyzed by one-way ANOVA with Dunnett's multiple comparisons test (ns $p$>0.05, *$p$<0.05, **$p$<0.01, ***$p \leq$ 0.001, ****$p \leq$ 0.0001). **C)-E)** Bulk RNA-seq of bryostatin- SC1 cells treated with 1 nM bryostatin for 72 hours. **C)** UMAP depicting FPKM (fragments per kilobase million) and TPM (transcripts per million) comparing DMSO control and bryostatin-treated cells. **D)** Volcano plot represents 154 significantly upregulated and 17 significantly downregulated genes (FDR<0.05, $log_2FC>2$). **E)** Heatmap for the 591 up and 204 down genes (FDR<0.05, total 795 genes) differentially expressed in DMSO versus Bryostatin-1 treated cells.

**Supplemental Table 1**: Antibodies used for flow cytometry

| Antibody, clone | Manufacturer | Catalog number |
|---|---|---|
| Human CD72-APC (Clone 3F3) | Biolegend | 316210 |
| Human CD72-PE (Clone 3F3) | Biolegend | 316208 |
| Human CD72-PE (Clone 3F3) | Invitrogen | MA1-19790 |
| Human CD72-BV421 (Clone J4-117) | BD Biosciences | 743794 |
| Human CD19-APC (Clone HIB19) | BD Biosciences | 555415 |
| Human CD19-PE (Clone HIB19) | Biolegend | 302208 |
| Human CD3-APC (Clone SK7) | Biolegend | 344812 |
| Human CD3-APCfire (Clone SK7) | Biolegend | 344840 |
| Human CD45-APC (Clone 2D1) | Biolegend | 368512 |
| Human CD45-PB (Clone 2D1) | Biolegend | 368540 |
| Human CD45-PE (Clone 2D1) | Biolegend | 368510 |
| Human CD22-FITC (Clone HIB22) | Biolegend | 302504 |
| Human CD22-APC (Clone HIB22) | Biolegend | 302510 |
| Human CD20 – BV421 (Clone L27) | BD Biosciences | 740002 |
| CD22 PE (Clone S-HCL-1) | BD Biosciences | 340708 |
| CD34 PerCP Cy5.5 (Clone 8G12), | BD Biosciences | 347213 |
| CD19 PE Cy-7 (Clone J3-119), | Beckman Coulter | IM1285U |
| CD10 APC (Clone HI10a), | BD Biosciences | 340922 |
| CD45 APC-H7 (Clone 2D1) | BD Biosciences | 560178 |
| CD38 BV510 (Clone HB-7) | Biolegend | 356612 |

| | | |
|---|---|---|
| Human CD107a-APC (Clone H4A3) | Biolegend | 328620 |
| Human CD69 – PE (Clone FN50) | Biolegend | 310906 |
| Human CD45RA-PE (Clone HI100) | Invitrogen | 12-0458-42 |
| Human CD62L-APC (Clone DREG-56) | BD Biosciences | 559772 |
| Human PD1-PB (Clone EH12.2H7) | Biolegend | 329916 |
| Human LAG3-PE (Clone T-47-530) | BD Biosciences | 565616 |
| Human TIM3-APC (Clone F38-2E2) | Invitrogen | 17-3109-42 |
